# Vacuum and sonication treatment enables efficient transient gene expression in various monocot and eudicot plant seedlings

**DOI:** 10.1101/2025.03.13.642903

**Authors:** Eugene Li, Yuan Geng, Tuyako R. Khristoforova, Yunqing Wang, Jolie W. Jones, Gozde S. Demirer

## Abstract

Transient gene expression in intact plants is essential for rapidly addressing biological questions, where the current toolkit can be improved for higher efficiencies and broader plant species range. Here, we introduce VAST (Vacuum and Sonication-Assisted Transient expression): a transient transformation method that substantially enhances gene expression efficiency, reproducibility, and versatility across diverse monocot and eudicot intact seedlings. By systematically optimizing plant growth conditions and incorporating vacuum infiltration and sonication pre-treatments prior to seedling co-culture with *Agrobacterium tumefaciens*, we significantly improved transient gene expression efficiency while minimizing tissue damage compared to existing methods in *Arabidopsis thaliana*. We further demonstrated the broad applicability of VAST by successfully transforming key model and crop species, including tomato, *Brassica rapa*, *Medicago sativa*, *Setaria italica* (foxtail millet), switchgrass, and wheat. We also demonstrated a case study using the VAST-mediated transient transformation, where a cross-species analyses of nitrate-responsive gene expression highlighted both the conserved and diverged biological responses between the two model crops, *A. thaliana* and *S. italica*. VAST’s simplicity, versatility, and efficiency provide a powerful tool for functional genomics, synthetic biology, and biotechnology research, opening new avenues for the rapid exploration of gene function, regulation, and editing in diverse plant systems.

## Introduction

Plants hold the key to addressing some of today’s most pressing global challenges, including food security, sustainability, and climate change. Plant genetic engineering offers a rapid and targeted path to producing sufficient, nutritious food while ensuring crop resilience under changing climate and mitigating the impact of agriculture on the environment. Moreover, non-food plants can serve as a scalable and sustainable source of biomanufacturing and bioenergy. However, plant genetic engineering is hampered by laborious, lengthy, and low efficiency plant transformation methods that are optimized for select species and cultivars.

Stable and heritable plant transformation is needed to answer many biological questions at the organismal level. Conversely, transient transformation typically manipulates somatic cells in intact plants without the aim of collecting progeny, which is highly useful for rapid study of gene function, protein localization, protein-protein or protein-DNA interactions, and various other synthetic biology and bioengineering applications. Moreover, transient transformation of gene editing machinery, such as CRISPR/Cas9, can be the first step to create transgene-free edited plants.^1,2^ Therefore, transient gene expression is a powerful capability with a potential to substantially accelerate plant genetic engineering.

Current methods and capabilities of transient transformation are limited. Protoplasts can be transfected in some species from some tissues, but these short-lived cells in solution – lacking cell wall, cellular identity, and tissue structure – are unsuitable for answering many biological questions. Another common method uses *Agrobacterium tumefaciens*, which is effective in leaves of a narrow range of eudicot species for transient experiments, such as *Nicotiana benthamiana*.^3^ Efforts to increase the efficiency and to extend these methods to other species have been limited, particularly in monocots that feed most of the world’s population. In 2014, Wu et al. developed the AGROBEST method for transient gene expression in *Arabidopsis thaliana* seedlings using *A. tumefaciens*,^4^ which was based on a prior method called FAST.^5^ In 2021, Nasti et al. extended this method to multiple solanaceous species using a method called Fast-TrACC.^6^ These methods co-incubate plant seedlings with *A. tumefaciens* in liquid culture to achieve transformation. Lastly, Zhang et al. developed a transient transformation method by infiltrating intact leaves with *A. tumefaciens* in *A. thaliana* and 7 other eudicot plant species.^7^ There are also approaches based on the use of viral vectors delivered via *A. tumefaciens.*^8^ Although these have been great additions to our current toolkit, the transient transformation efficiency can be further enhanced and currently no method exists for routine transient transformation of monocots. Therefore, there is still a critical need for efficient transient transformation methods in intact eudicot and monocot plants.

Here, we present VAST (<u>V</u>acuum <u>a</u>nd <u>S</u>onication-assisted <u>T</u>ransient expression); a transient gene expression tool that significantly improves transformation efficiency, reproducibility, and species range across both eudicot and monocot intact seedlings. By incorporating vacuum and sonication pre-treatments before *A. tumefaciens* co-cultivation, VAST expands transformation capabilities without causing substantial tissue damage. We systematically optimized transformation protocols across multiple species, documented the optimization process using fluorescent reporter genes and quantitative real-time PCR, and accounted for plant health with mock and treatment samples. This study represents a successful demonstration of transient gene expression in intact monocot seedlings and is a major step forward in broadening the transient transformation toolkit in other eudicot plant species. Lastly, to illustrate the feasibility and impact of this tool, we present a case study comparing nitrate availability responses in two model plants: the eudicot *A. thaliana* and the monocot *S. italica*.

## Results

### Improving transient expression efficiency and uniformity in Arabidopsis thaliana leaves through vacuum and sonication treatments (VAST)

We sought to develop a transient expression system with high efficiency, uniformity, and broader species range by optimizing plant growth conditions, treatments at the beginning of co-cultivation with *A. tumefaciens*, and co-cultivation conditions. For this, 5-day-old *A. thaliana* Col-0 wild-type (WT) seedlings were transformed with the *A*. *tumefaciens* GV3101 strain carrying a yellow fluorescent protein (YPET) expression plasmid (**Supplementary Table 1** for plasmid details). At 4 days post-infection (dpi), YPET expression was imaged with a fluorescence microscope, and *YPET* relative gene expression levels were measured via quantitative real-time PCR (qPCR) to compare efficiencies of different conditions and treatments.

We first tested two plant growth conditions. Apart from a 2-day liquid co-culture with the *A*. *tumefaciens* suspension, *A. thaliana* seedlings were germinated and maintained throughout the experiment either in liquid Murashige and Skoog (MS) plant growth media, as in previous methods of AGROBEST^4^ and Fast-TrACC^6^, or on solid MS media. Seedlings grew faster and were substantially healthier on solid MS compared to those in liquid MS, although *YPET* expression levels were comparable under both conditions without any additional treatment (**Figure 1A, B**).

**Figure 1.**
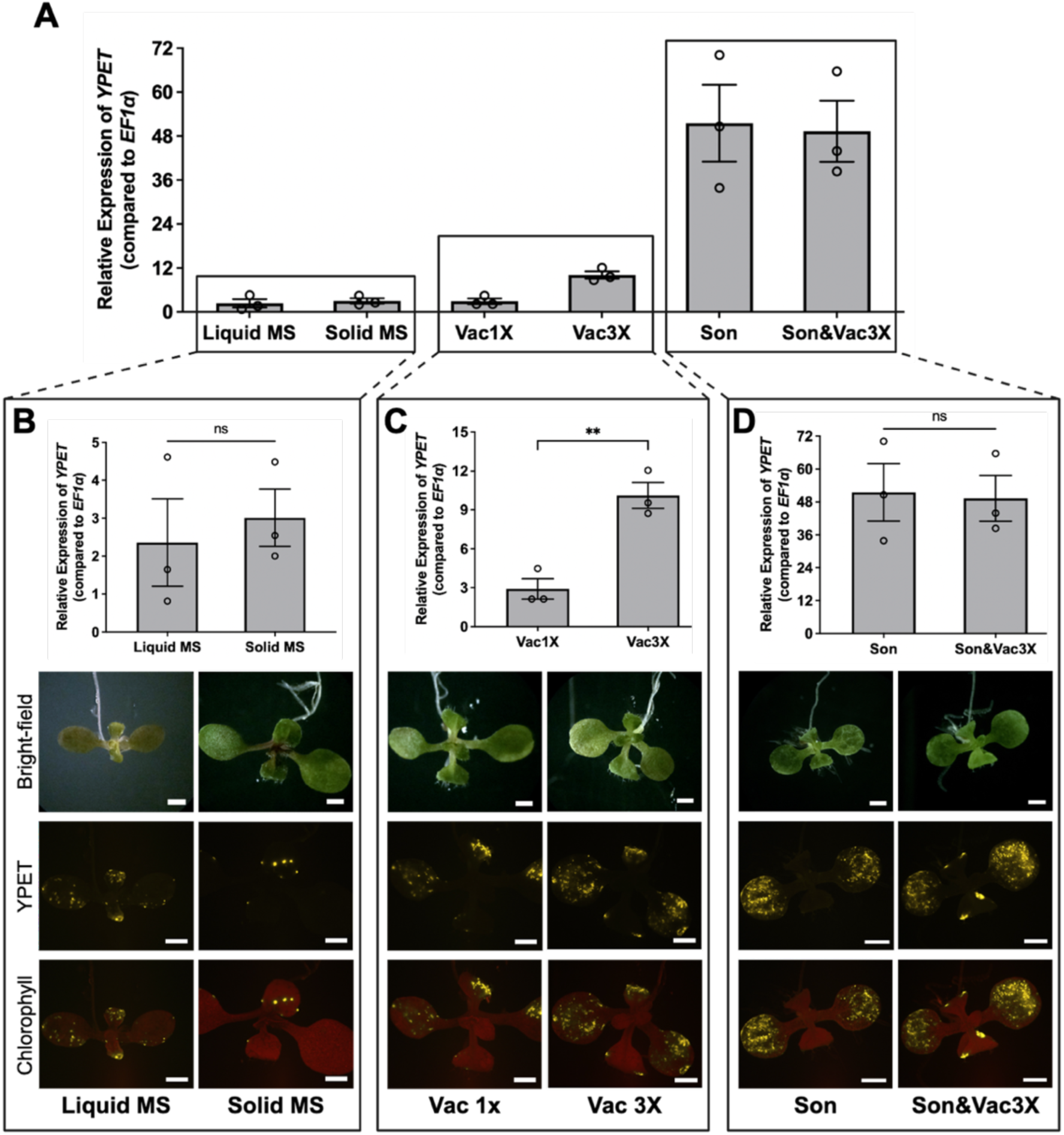
Optimization of transient transformation conditions in *A. thaliana* seedlings. **(A)** Relative expression of *YPET* in *A. thaliana* seedlings with different treatment conditions, obtained via quantitative PCR (qPCR) and normalized to the reference gene *EF1α*. Bars: mean ± standard error of the mean (SEM), N = 3 biological replicates, each replicate containing 5 pooled seedlings. **(B-D)** Relative expression of *YPET* and representative fluorescence microscopy images of seedlings treated with different conditions (note the different y-axis scale in each plot): liquid Murashige and Skoog (MS) media (Liquid MS) versus solid MS media (Solid MS) **(B)**; vacuum once (Vac 1x) versus vacuum three times (Vac 3x) **(C)**; sonication (Son) versus combined sonication and vacuum three times (Son&Vac3x) **(D)**. Statistical analysis was performed using a student’s *t*-test. \*\**p* < 0.01. ns: not significant. The experiment was independently repeated three times with similar results. Channels: Bright-field, YPET (yellow), and Chlorophyll (red) overlaid with YPET. All scale bars: 1 mm.

Next, seedlings germinated on solid media were transferred into an *A. tumefaciens* suspension and subjected to the following physical treatments to increase tissue penetration of *A. tumefaciens* and improve expression efficiency and uniformity: vacuum once (Vac1x), vacuum 3 times (Vac3x), bath sonication for 20 seconds (Son), or 20 seconds sonication combined with 3 cycles of vacuum (Son&Vac3x) (**Figure 1A**). Subsequently, the seedlings were co-cultured with *A. tumefaciens* for two days and transferred to solid MS media until imaging and qPCR analyses.

Fluorescence imaging showed that vacuum application increased expression uniformity, where *YPET* expression levels were more than 3.3 times higher in Vac3x compared to Vac1x and solid MS no treatment (**Figure 1A, C**). Sonication treatment also significantly enhanced transformation efficiency and uniformity as seen in fluorescence images (**Figure 1D**), with a 17-fold increase in *YPET* expression compared to no treatment (solid MS) and 5-fold increase compared to Vac3x (**Figure 1A**). While the combined sonication and vacuum treatment did not further increase *YPET* expression as quantified by qPCR in average of 15 seedlings, it did promote expression both in cotyledons and true leaves (**Figure 1D**). In addition to the one set of representative fluorescence images in **Figure 1**, three more sets of images per treatment from different biological replicates are provided in **Supplementary Figure 1**, showing the reproducibility of transformations.

We next examined the effect of *A. tumefaciens* concentration used in 2-day liquid co-culture on the transient gene expression efficiency. Increasing *A. tumefaciens* concentration from OD_600_ 0.14 to 0.3 or 0.5 did not result in further improvement of *YPET* expression in *A. thaliana* seedlings as shown by fluorescence imaging and qPCR (**Supplementary Figure 2**). We also evaluated the effects of varying vacuum infiltration durations (2, 5, and 8 minutes) and sonication times (10, 20, and 40 seconds) on transformation efficiency and stress response (**Supplementary Figure 3**). Based on fluorescence imaging, the combination of 5 minutes of vacuum and 20 seconds of sonication produced the strongest and most uniform YPET fluorescence signal (**Supplementary Figure 3A, B**), while also minimizing the stress response, as quantified via the expression levels of a canonical stress gene, Pathogenesis-Related protein 1 *(PR1)* (**Supplementary Figure 3C, D**). qPCR analysis revealed no statistically significant differences in YPET expression across these conditions (**Supplementary Figure 3C, D**). In conclusion, given imaging results, we recommend 5 minutes of vacuum and 20 seconds of sonication conditions as the optimized parameters for VAST.

Next, we validated that the expression is originating from plant cells, and not in *A. tumefaciens,* through the use of an RNA-to-cDNA conversion kit for qPCR that is selective for eukaryotic mRNA (see methods), and by expressing a YPET gene interrupted with the second intron of the ST-LS1 gene from potato^9^. In both cases, we detected the plant-originated YPET expression in fluorescence microscopy imaging (**Supplementary Figure 4**) and qPCR (**Figure 1A**). Lastly, we demonstrated that VAST transformation is also effective in non-sterile and soil-grown Arabidopsis plants (**Supplementary Figure 5**), in addition to those grown on plates, thereby broadening the applicability of the system.

In summary, by modifying the plant growth medium from liquid to solid and introducing and systematically optimizing vacuum and sonication pre-treatments, we significantly improved the efficiency, uniformity, and reproducibility of transient gene expression in *A. thaliana* seedlings. We named this method VAST – vacuum and sonication-assisted transient expression.

### VAST is more efficient than previous methods and reduces cellular stress

We compared the efficiency of optimized VAST with that of other transient gene expression tools. For this, the same YPET construct was transformed into Col-0 *A. thaliana* seedlings following the published protocols of AGROBEST^4^ and Fast-TrACC^6^, and our optimized VAST protocol (**Figure 2A**). 4-dpi fluorescence imaging revealed strong and uniform *YPET* expression in cotyledon and true leaves of VAST-transformed seedlings, whereas AGROBEST and Fast-TrACC-transformed seedlings of the same age showed lower and more sporadic expression (**Figure 2B**). When quantified with qPCR for YPET expression using 20 seedlings in each condition, VAST achieved an average of 25-and 10-fold higher expression than AGROBEST and Fast-TrACC, respectively (**Figures 2C**). In addition to the one set of representative fluorescence images in **Figure 2B**, three more sets of images from different biological replicates are provided in **Supplementary Figure 6.**

**Figure 2.**
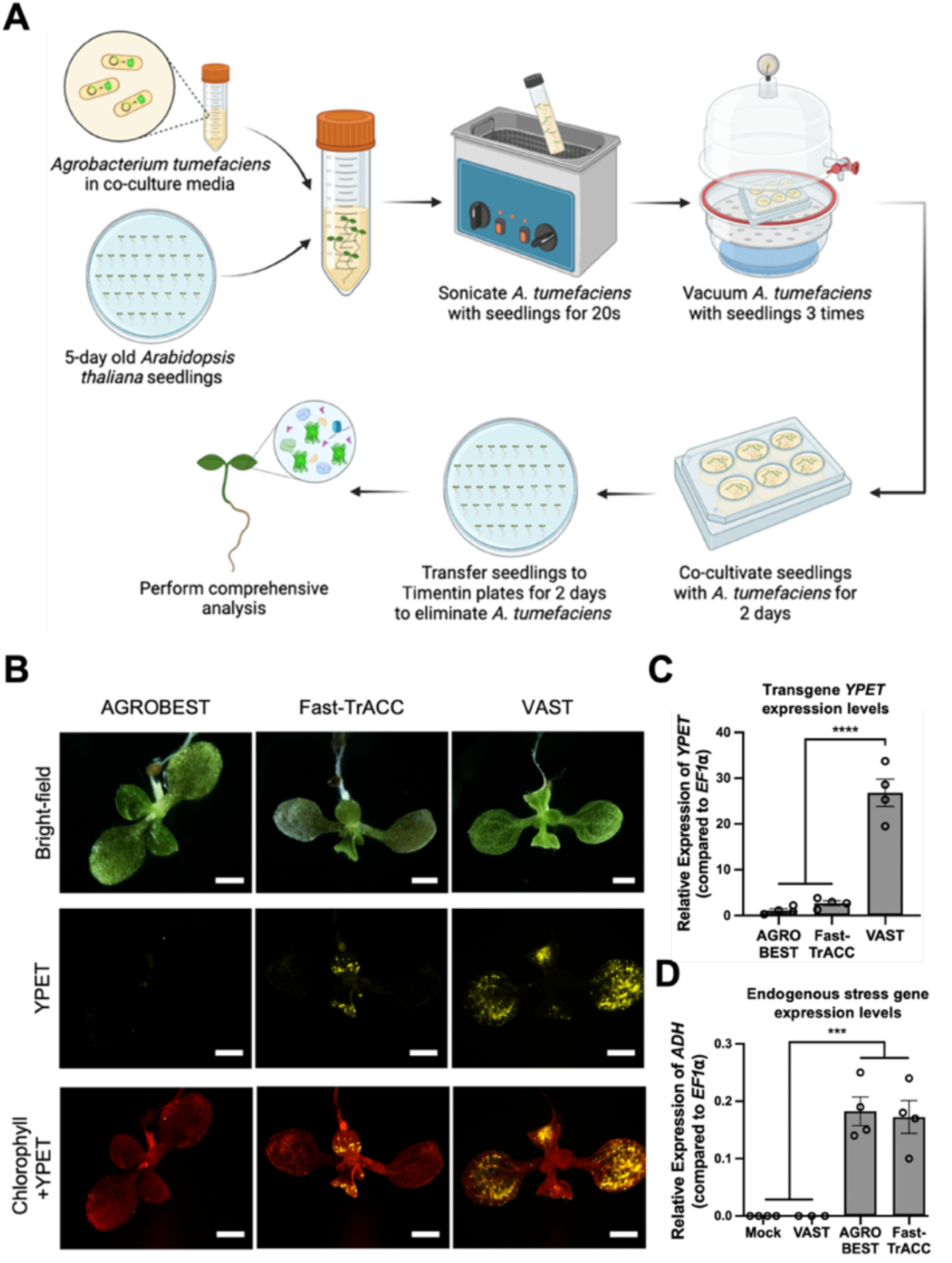
Comparison of VAST with other transient expression methods in *A. thaliana* using GV3101. **(A)** Schematic workflow of VAST. **(B)** Representative fluorescence microscopy images of *A. thaliana* seedlings at 4-dpi. All scale bars: 1 mm. Channels: Bright-field, YPET (yellow), and Chlorophyll (red) overlaid with YPET. **(C)** qPCR analysis of *YPET* expression at 4-dpi for AGROBEST, Fast-TrACC, and VAST, normalized to *EF1α*. **(D)** qPCR analysis of the hypoxia-induced stress gene *ADH* at 4-dpi, normalized to *EF1α*. For **(C-D)**, data represent mean ± standard error from 4 biological replicates with 5 seedlings per replicate. The experiment was independently repeated three times with similar results. Statistical outliers were identified and excluded using Grubbs’ test. One-way ANOVA with Tukey’s test was used for analysis, with significance indicated as ****p < 0.0001, ***p < 0.0002, **p < 0.0021, *p < 0.0332.

In addition to the GV3101 strain used in above experiments, we also tested another commonly used *A. tumefaciens* strain, C58C1(pTiB6S3ΔT)^H^. C58C1 achieved more efficient expression overall, compared to GV3101, especially using AGROBEST and Fast-TrACC. However, VAST still significantly outperformed both methods as seen in fluorescence images (**Figure 3A**) and qPCR quantification of YPET gene expression (**Figure 3B**). It is also apparent from brightfield images that VAST-transformed seedlings are substantially healthier than others (**Figure 3A**).

**Figure 3.**
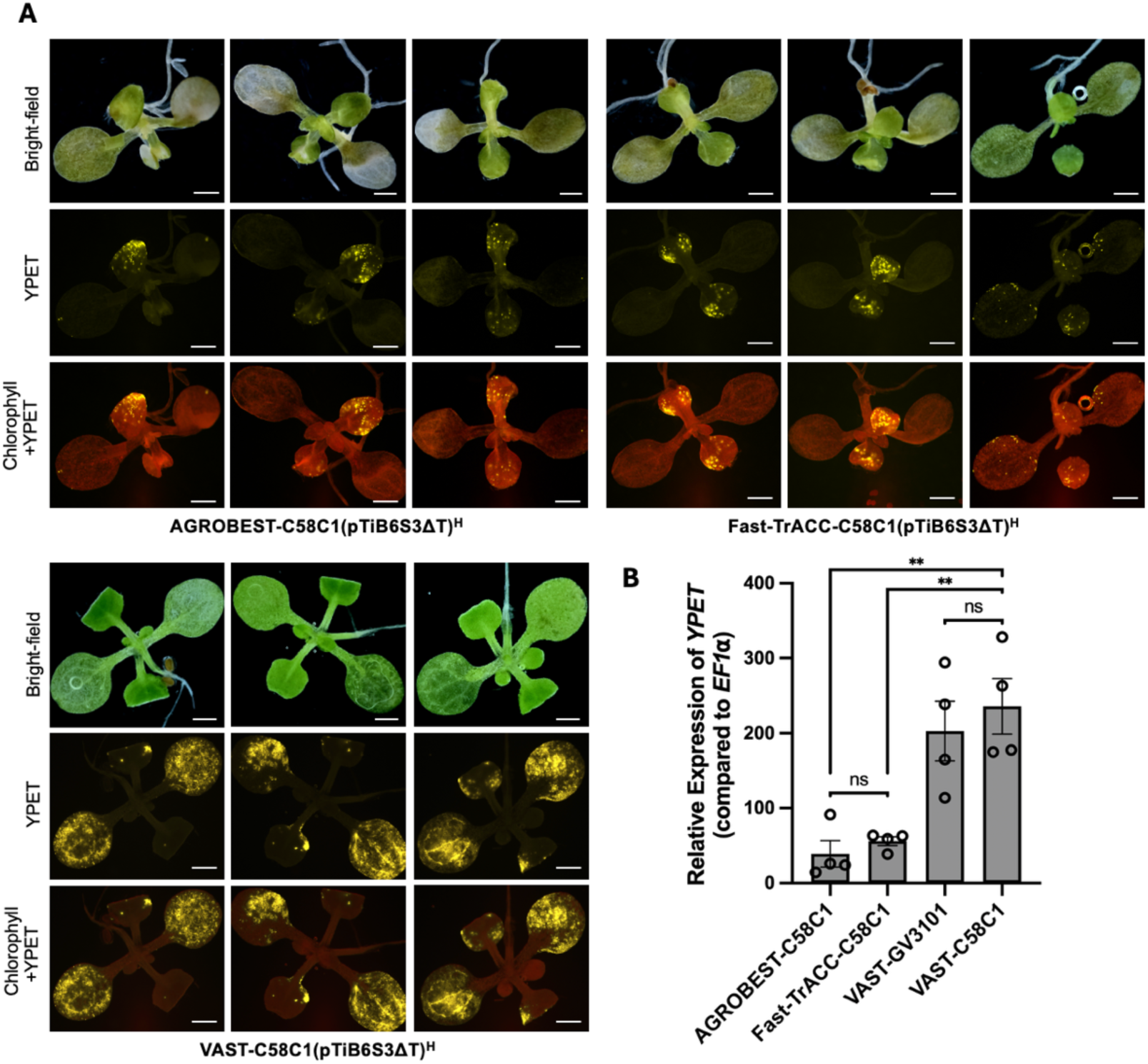
Comparison of VAST with other transient expression methods in *A. thaliana* using C58C1(pTiB6S3ΔT)^H^. **(A)** Representative fluorescence microscopy images of *A. thaliana* seedlings transformed using AGROBEST, Fast-TrACC, or VAST with *A. tumefaciens* strain C58C1(pTiB6S3ΔT)^H^. Channels: bright-field (top), YPET fluorescence (middle), and merged YPET + chlorophyll (bottom). Scale bars = 1 mm. **(B)** qPCR analysis of YPET expression at 4-dpi, normalized to the housekeeping gene EF1α. Data represent mean ± standard error from 4 biological replicates (5 seedlings per replicate) from a representative experiment. The experiment was independently repeated three times with similar results. Statistical outliers were identified and excluded using Grubbs’ test. One-way ANOVA with Tukey’s test was used for statistical analysis. Significance levels: **p < 0.0021, ns = not significant.

Seedlings transformed with AGROBEST and Fast-TrACC for YPET expression exhibited more leaf tissue damage compared to seedlings transformed with VAST or mock treatment as seen in five representative brightfield images in **Supplementary Figure 7**. We hypothesized that this might result from prolonged incubation in liquid MS, both pre- and post-transformation, leading to hypoxia-induced stress and damage. To assess this hypothesis, we measured the endogenous expression levels of alcohol dehydrogenase (*ADH*) in YPET transformed seedlings. ADH is a critical enzyme in anaerobic metabolism that supports ATP production under hypoxic conditions^10^. Elevated *ADH* expression serves as a marker for hypoxic stress^11^. Using qPCR, we measured *ADH* expression in seedlings transformed with VAST, AGROBEST, Fast-TrACC, and a mock treatment, which consists of seedlings grown on solid media, incubated in liquid co-culture without *A. tumefaciens* for 2 days, and transferred to solid media post-incubation **(Supplementary Figure 6D).** Our results showed that AGROBEST- and Fast-TrACC-transformed seedlings exhibited *ADH* stress gene expression levels 103- and 97-fold higher than the mock treatment at 4-dpi, respectively, confirming substantial hypoxia stress. In contrast, VAST-transformed seedlings showed *ADH* levels similar to the mock treatment (**Figure 2D**).

In addition to the hypoxia-induced stress, we also measured general oxidative stress levels using the DAB (3, 3’-diaminobenzidine) histological staining, where DAB reacts with hydrogen peroxide (H₂O₂) in the presence of endogenous peroxidases to produce a brown precipitate, which localizes at sites of H₂O₂ accumulation. Results showed minimal ROS accumulation in VAST-transformed seedlings compared to the mock, whereas substantial browning and significant oxidative stress was observed in AGROBEST and Fast-TrACC-transformed seedlings (**Supplementary** Figure 8A). Quantification of the DAB staining brown color intensity corroborates this trend, in which the small increase in VAST samples is not statistically significant, while AGROBEST and Fast-TrACC samples have 24-times more DAB staining compared to the mock treatment (**Supplementary Figure 8B**).

To assess the persistence of gene expression and long-term plant health, we extended *YPET* and *ADH* expression observations to 10-dpi. Seedlings transformed with AGROBEST and Fast-TrACC displayed almost no *YPET* expression at 10-dpi (**Supplementary Figure 9A, B**). Even though the *YPET* expression in VAST-transformed seedlings were lowered due to the emergence of new leaves and expansion of old leaves, there was still substantial *YPET* signal remaining in VAST-transformed seedlings at 10-dpi, with 133- and 10-fold increase compared to the AGROBEST and Fast-TrACC, respectively (**Supplementary Figure 9C, D**). Moreover, seedlings transformed with AGROBEST and Fast-TrACC were fragile at 10-dpi, had white cotyledons, and were disassembling even with gentle handling. This unhealthy plant phenotype was corroborated by the stress gene *ADH* expression levels of 35- and 49-fold higher than of mock-treated plants, respectively, for AGROBEST and Fast-TrACC-transformed plants. In contrast, VAST-transformed seedlings maintained relatively healthy growth and *ADH* levels similar to mock-treated plants, indicating no hypoxia-induced damage (**Supplementary Figure 9D**).

Next, we conducted further in-depth characterization of the temporal and spatial performance of the VAST-mediated expression in *A. thaliana*. At 4-dpi, the expression is very strong and covers almost the entire leaf area, which gets sparser at 10-dpi given leaf growth but still strong and easily detectable. By 16-dpi, the leaves are significantly larger and there are many new leaves, challenging the detection of expression (**Supplementary Figure 10**). Therefore, the expression can be monitored in adult plants until 3-weeks (21-days) age. In terms of spatial properties within the transformed *A. thaliana* leaf, strong expression is obtained both in the epidermal and mesophyll cell layers as seen in the confocal microscopy images, regardless of the *Agrobacteria* strain used (**Supplementary Figure 11, 12A**). Detailed analysis of YPET expression at the single-cell resolution revealed transformation efficiencies up to 90% in some *A. thaliana* seedlings with an average of 60% in all transformed seedlings (**Supplementary Figure 11D, E**).

### Efficient transient transformation of various eudicot plant species with VAST

Next, we expanded the VAST transient transformation method to other eudicot plant species. Seedlings of *Solanum lycopersicum* (Micro-Tom), *Brassica rapa* (Ames 30080)*, Medicago sativa* (Bulldog 805) were transformed with the *A*. *tumefaciens* strain GV3101 carrying the same *YPET* expression construct to evaluate and optimize the transient gene expression efficiency of VAST in a broader species range. These eudicot plant species were selected given both their importance as model research organisms and their potential impact on many food and environmental applications.

The VAST protocol established in *A. thaliana* was used as a starting point in these species with some further optimizations (see **Supplementary Table 2** and Methods for details). For instance, *A. tumefaciens* concentration was increased from OD_600_ 0.14 to 0.4 for all three species. For *S*. *lycopersicum* and *M. sativa*, the sonication duration was extended from 20 seconds to 30 and 45 seconds, respectively. Since *B. rapa* seedlings were very sensitive to the *A. tumefaciens* infection, the sonication duration was shortened to 15 seconds and the co-cultivation duration was reduced from 2 days to 1 day to minimize the tissue damage caused by *A. tumefaciens* infection in *B. rapa*.

Using these optimized VAST transformation settings in *S*. *lycopersicum*, *B*. *rapa*, and *M*. *sativa*, we observed strong *YPET* expression in both cotyledons and true leaves of all three species at 4-dpi using fluorescence microscopy (**Figure 4A**). We further checked the *YPET* expression at 5-dpi using confocal microscopy, which corroborated strong and uniform reporter expression in all species. Confocal fluorescence microscopy also showed that the YPET protein was localized both in nucleus and cytoplasm of transformed cells (**Figure 4B**), as expected from a small fluorescent protein without a subcellular tag. The nuclear localization validates that the fluorescence signal indeed belongs to YPET instead of being background, autofluorescence, or tissue damage-related signal. In addition to the one set of representative fluorescence and confocal microscopy images for each species in **Figure 4**, three more sets of transformed and mock seedling (negative control) fluorescence microscopy images of all species from different biological replicates are provided in **Supplementary Figure 13.**

**Figure 4.**
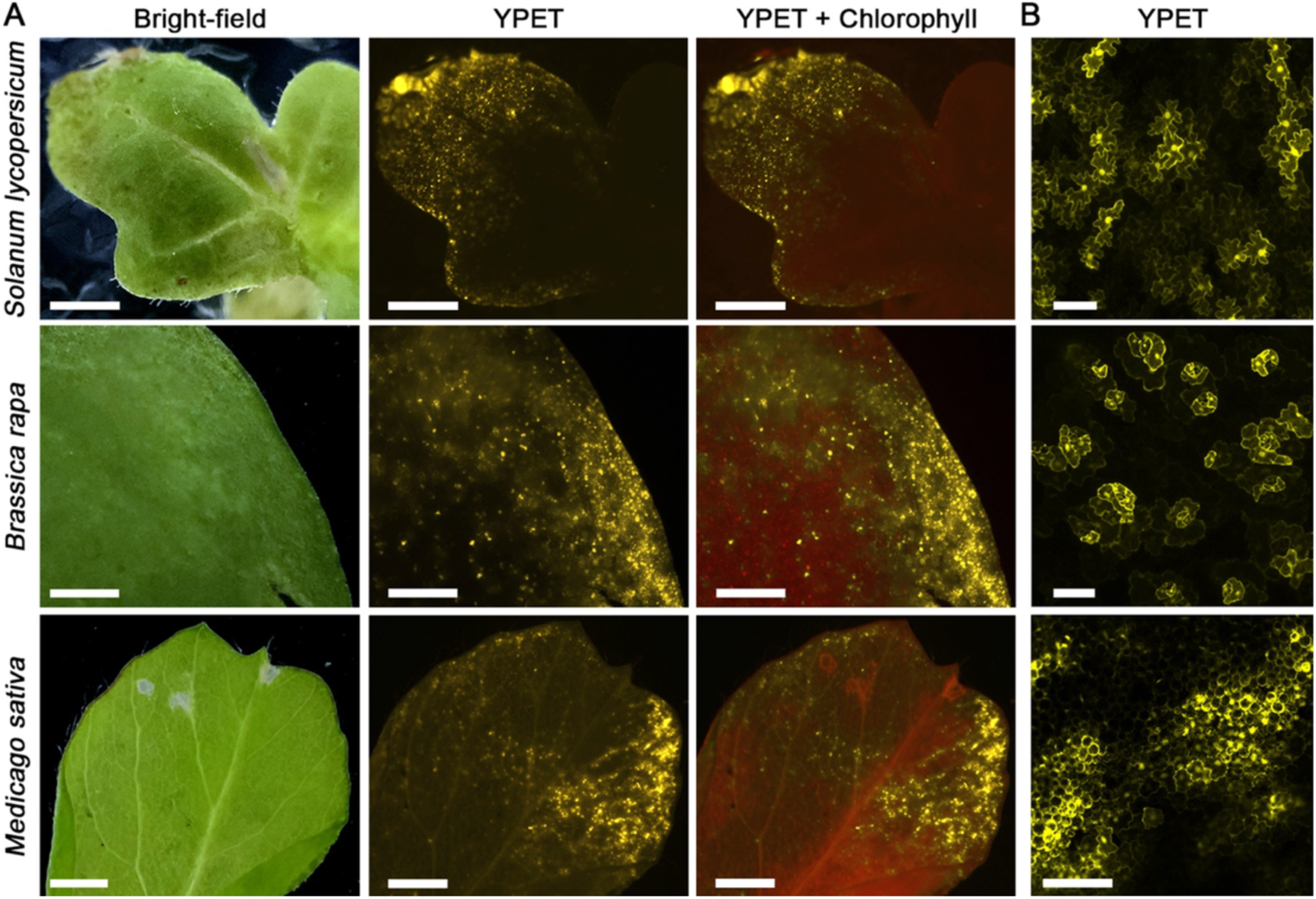
VAST-mediated transient transformation of other eudicot plant species. **(A)** Representative fluorescence microscopy images of VAST-transformed *S*. *lycopersicum*, *B*. *rapa*, and *M*. *sativa* leaves at 4-dpi. Channels: Bright-field, YPET (yellow), and Chlorophyll (red) overlaid with YPET. All scale bars, 1 mm. **(B)** Representative confocal fluorescence microscopy images of the same seedlings expressing *YPET* at 5-dpi. All scale bars, 100 µm.

### Efficient transient transformation of various monocot plant species with VAST

We next optimized the VAST-mediated transient gene expression in several important monocot species: *Setaria italica* (foxtail millet), *Panicum virgatum* (switchgrass), and *Triticum aestivum* (wheat). Foxtail millet is a C4 annual grass that is resistant to drought, low soil fertility, and high temperatures, and contains high levels of calories, nutrients, protein, and fiber. Switchgrass is one of the primary crops for bioenergy production, and wheat is a staple global food source. Compared to eudicots, monocots are less susceptible to *A. tumefaciens* infection, making their genetic manipulation challenging, with no current tools for transient transformation.

In our initial experiments, use of the optimized VAST protocol from eudicots did not yield sufficient gene expression in these monocot species, and hence, we performed further rounds of optimization. To enhance transformation efficiency of VAST in monocots, *A. tumefaciens* AGL1 strain was used instead of GV3101 to deliver the YPET expression cassette. Additionally, *A. tumefaciens* concentration was increased to OD_600_ = 0.4 from 0.14, and the sonication duration was extended to 30 seconds followed by 4 rounds of vacuum treatment.

Using this optimized protocol, we observed YPET expression in the leaves of all three monocot species tested at 4-5 dpi (**Figure 5A**). Across multiple rounds of independent transformation events, *S*. *italica* has shown the strongest and most uniform expression compared to other monocots. Expression in switchgrass was strong but more sporadic, typically existing only in certain parts of the transformed leaves. Compared to *S*. *italica* and switchgrass, wheat showed lower transformation efficiency, where in each transformed seedling, only a small number of cells expressed YPET (**Figure 5A**). VAST-mediated transient gene expression in monocots was also verified in all species using confocal fluorescence microscopy imaging, where YPET expression localized to both the cytosol and nuclei of transformed cells as expected (**Figure 5B**). Fluorescence microscopy images of mock-transformed *S*. *italica* and switchgrass seedlings are provided in **Supplementary Figure 14**.

**Figure 5.**
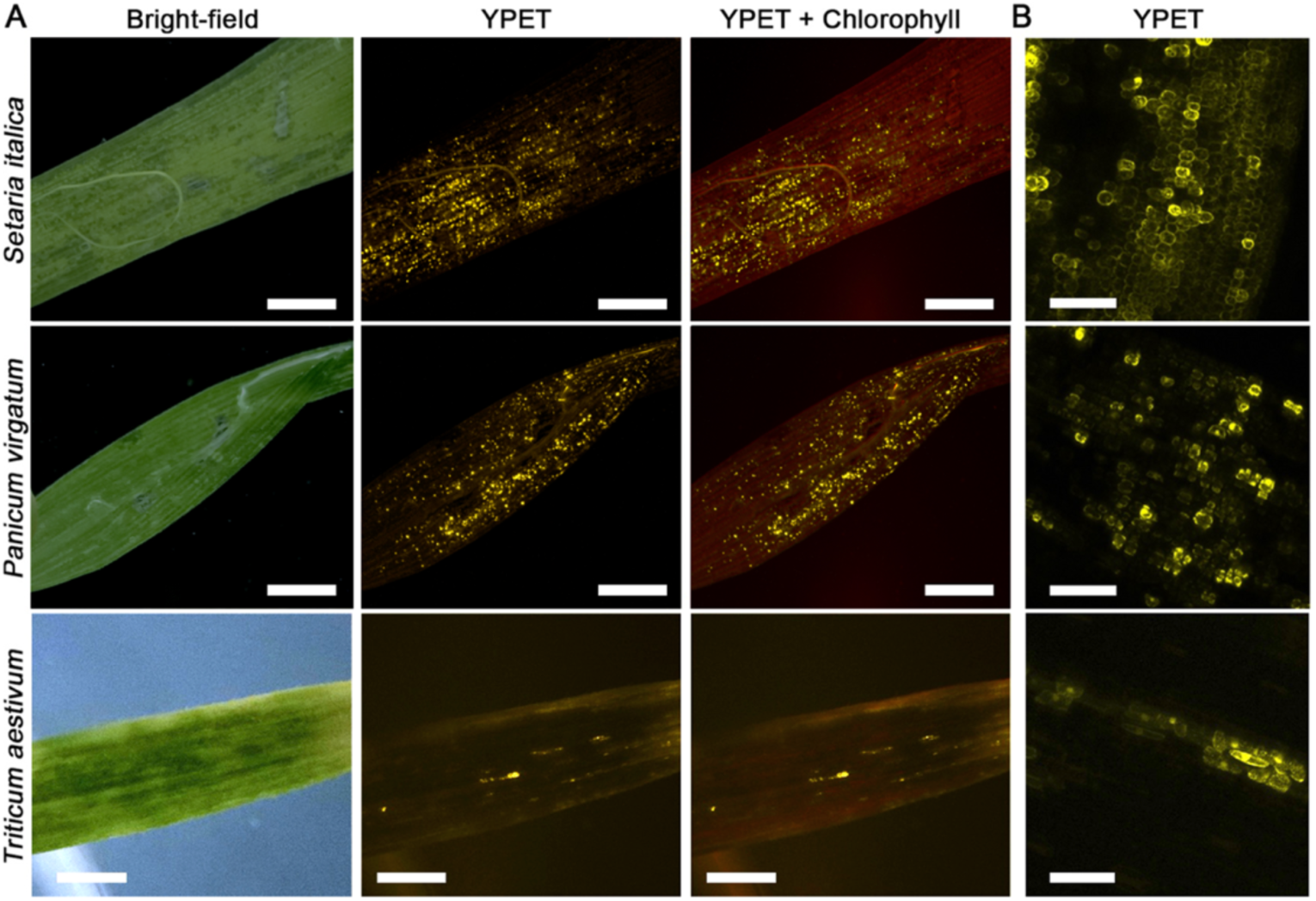
VAST-mediated transient transformation of monocot plant species. **(A)** The fluorescence microscopy images of VAST-transformed *S*. *italica*, *P*. *virgatum*, and *T*. *aestivum* leaves at 4-dpi. Scale bars, 1 mm. **(B)** Confocal microscopy images of the same seedlings expressing YPET at 5-dpi. Scale bars, 100 µm. Channels: Bright-field, YPET (yellow), and Chlorophyll (red) overlaid with YPET.

In addition to the AGL1 strain, we evaluated another commonly used *A. tumefaciens* strain in monocots, EHA105. While EHA105 did enable transformation in *S*. *italica*, YPET expression was noticeably weaker and sparsely distributed as seen both in fluorescence and confocal microscopy images (**Supplementary Figure 15A, B**). This observation was supported by qPCR analysis, which showed 16-fold higher YPET expression in AGL1-transformed seedlings compared to those transformed with EHA105 (**Supplementary Figure 15C**). Moreover, leaf tissue from EHA105-treated samples exhibited more visible damage, as indicated by increased browning. This was accompanied by slightly higher expression of a representative stress marker gene, *catalase*, as measured by qPCR, although the difference was not statistically significant (**Supplementary Figure 15D**). Catalase gene expression levels that were not significantly elevated indicate that the VAST treatment, especially when used with the AGL1 strain for monocots, does not cause major oxidative stress or damage (**Supplementary Figure 15D**).

In terms of spatial properties within a transformed *S. italica* leaf, strong expression is obtained both in the epidermal and mesophyll cell layers as seen in the confocal microscopy images (**Supplementary Figure 12B, 15B**). Detailed analysis of reporter expression at the single-cell resolution revealed transformation efficiencies up to 40% in some seedlings, with an average of 23% in over 20 seedlings (**Supplementary Figure 15E**).

Overall, although the efficiency varies among different monocot species, our results suggest that VAST can be used to achieve transient gene expression across a broad range of plant species, including monocots, all of which currently lack rapid transient transformation capabilities.

### Functional analysis of plant nitrate response via VAST transient transformation

VAST’s efficiency for rapid delivery of gene constructs into intact seedlings makes it a valuable tool for studying gene and promoter function *in planta*. It enables answering physiological questions without needing stable transformants or questions that would not be possible to address with current transient transformation tools, such as protoplast transfection or *Agroinfiltratio*n of *N. benthamiana* leaves. To illustrate this, we performed a case study characterizing and comparing the nitrate response of the model eudicot *A. thaliana* and model monocot *S*. *italica* by VAST-transforming seedlings with a nitrate-sensing circuit. Besides our method, the only way to efficiently and transiently transform these species is to use protoplasts, which do not live long enough to complete all aspects of this study, nor they could provide a relevant physiological response for nitrate availability that relies on systemic response. Hence, without our method, this study would have only been possible via time-intensive stable transformation process.

The transformed circuit employs a Nitrate-Regulated Promoter (NRP)^12^, a synthetic promoter activated by nitrate, which has been used in prior studies to assess the wild type and engineered tomato response to nitrogen availability.^13^ Our construct was designed with two components **(Figure 6A)**: the first component places the luciferase GeNL^14^ reporter gene under the control of NRP, allowing the nitrate-induced promoter activity to be visualized and quantified in a high-throughput manner using a plate reader. The second component includes NLuc luciferase driven by a constitutive promoter, providing an internal standard to normalize for cell count, health, and transformation efficiency, ensuring accurate, quantitative, and ratiometric assessment of NRP’s activity, hence plant’s response to nitrate availability.

**Figure 6.**
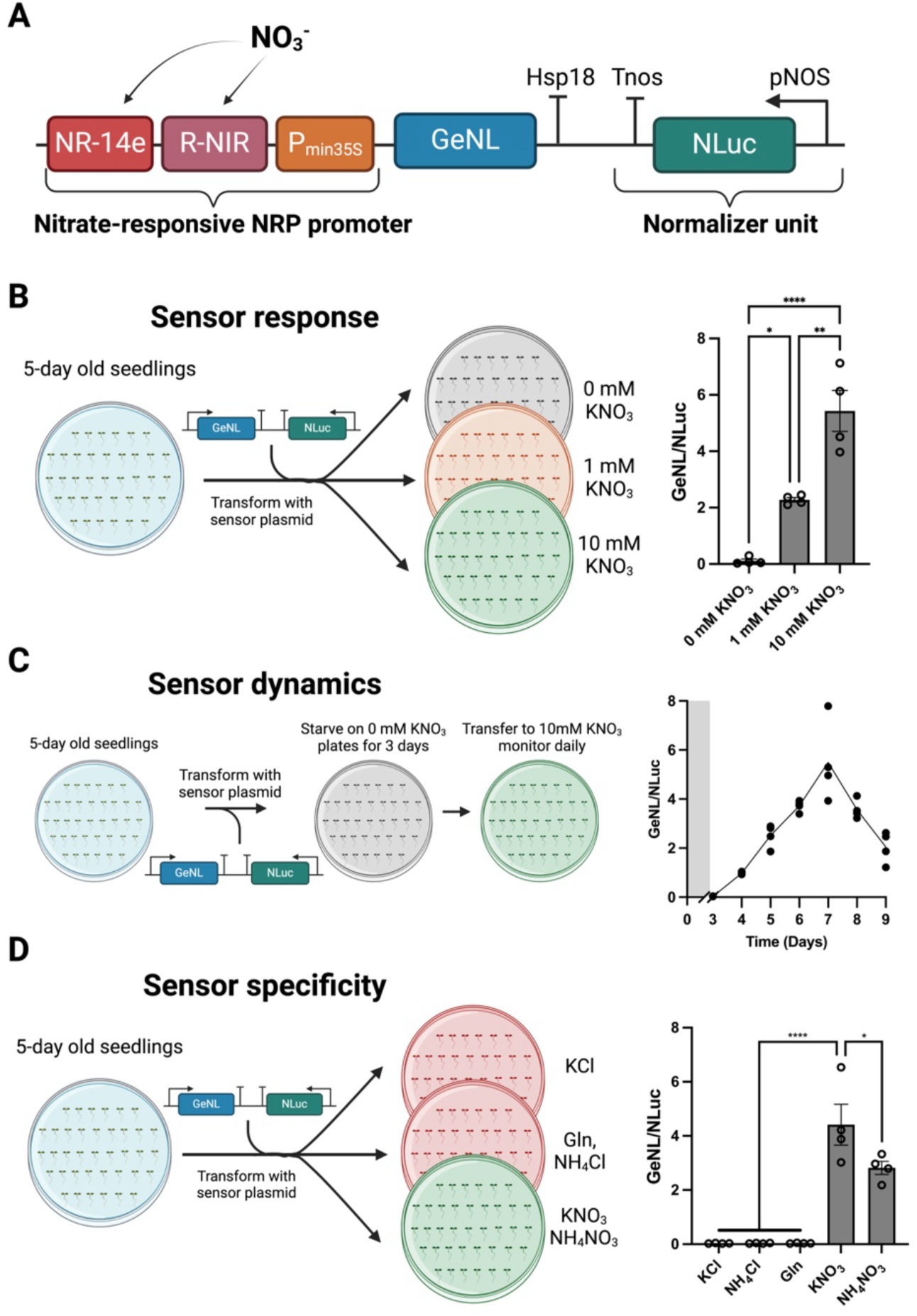
Functional analysis of the Nitrate-Regulated Promoter (NRP) in *A. thaliana.* **(A)** Schematics of the constructs: GeNL (Green Enhanced Nano-lantern), NLuc (NanoLuc luciferase), pNOS (nopaline synthase promoter), Tnos (nopaline synthase terminator), and Hsp18 (heat shock protein 18.2 terminator) (**Supplementary Table 1**). **(B)** Workflow schematic for testing sensor response to 0-, 1-, and 10-mM nitrate. Data represent mean ± standard error from 4 biological replicates (5 seedlings per replicate). Statistical outliers were identified and excluded using Grubbs’ test. One-way ANOVA with Tukey’s test was used: ****P< 0.0001, ***P< 0.0002, **P< 0.0021, *P< 0.0332. **(C)** Workflow schematic for testing sensor dynamics after transferring plants to 10 mM nitrate plates. Each dot represents a biological replicate (4 biological replicates with 5 seedlings per replicate), and the line represents the mean of the replicates. The gray-shaded area indicates the nitrogen starvation period. **(D)** Workflow schematic for testing sensor specificity in response to various nitrogen sources or KCl at 10 mM. Gln: glutamine. Data represent mean ± standard error from 4 biological replicates (5 seedlings per replicate). One-way ANOVA with Tukey’s test was used: ****P< 0.0001, ***P< 0.0002, **P< 0.0021, *P< 0.0332.

To first test whether the VAST-transformation of this construct results in a similar NRP response in *A. thaliana* and then to assess the operational range of NRP activity, VAST-transformed *A. thaliana* seedlings were transferred to ½ MS plates containing 0-, 1-, or 10-mM nitrate for three days, followed by luminescence measurement **(Figure 6B)**. Similar to prior studies in tomato, NRP activity increased with nitrate concentration, demonstrating its functionality within a 0-10 mM range, which is representative of nitrate levels typically found in agricultural soils. These findings confirm the ability of VAST to replicate previous transformation outcomes and NRP’s suitability for detecting nitrate in plant systems under agronomically relevant conditions^13,15^.

Next, we examined the temporal dynamics of NRP activity and nitrate response in VAST-transformed seedlings. Transformed *A. thaliana* seedlings were grown on nitrogen-free ½ MS plates for three days to reduce NRP activity to baseline levels, then transferred to ½ MS plates containing 10 mM nitrate **(Figure 6C)**. Daily monitoring of luciferase activity revealed a sustained rise in NRP activity over four days as nitrate accumulated in leaf tissues. Following this period, luciferase activity declined, potentially because as the seedlings grew, nitrogen from older transformed leaves has been recycled into newer leaves. Temporal dynamics investigation of *A. thaliana* nitrate response performed here would not have been possible to study in protoplasts.

Finally, we assessed the specificity of NRP for nitrate over other nutrients or nitrogen sources in *A. thaliana*. Transformed seedlings were placed on plates containing various nitrogen sources or nitrogen-free plates for three days, followed by luminescence measurement **(Figure 6D).** NRP was specifically activated by nitrate and showed no activation in response to other nitrogen sources, such as ammonium or glutamine, or to KCl. In addition, the presence of ammonium suppressed NRP activity, consistent with previous findings^12^.

To compare the nitrate availability responses of *A. thaliana* to that of a monocot, the same experiment was performed in *Setaria italica*, a monocot model species closely related to maize and sorghum **(Supplementary Figure 16).** This comparison, enabled by VAST transformation, revealed critical information on the physiological and developmental differences between these two species. Similar to the results in *A. thaliana*, NRP activity in *S. italica* increased in response to nitrate, confirming its functionality in monocots even though the regulatory elements in the NRP promoter originate from *A. thaliana*. This hints a conservation between some nitrogen regulatory elements in monocots and eudicots. However, *S. italica* responded to nitrate availability and reverted to baseline significantly quicker than *A. thaliana* (4 vs 7 days), potentially suggesting a different nitrogen use efficiency or amount needed in *S. italica* compared to *A. thaliana*.

## Discussion

In this study, we developed VAST (Vacuum and Sonication-assisted Transient expression), an improved transient transformation method that significantly enhances gene expression efficiency, reproducibility, and applicability in various intact monocot and eudicot seedlings, such as *Arabidopsis thaliana*, tomato, *Brassica rapa*, *Medicago sativa*, *Setaria italica*, switchgrass, and wheat. By systematically optimizing physical treatments of vacuum infiltration and sonication, and refining plant growth conditions (**Figure 1, Supplementary Figure 3**), VAST achieves substantial improvement over existing transient expression tools. Specifically in *Arabidopsis thaliana*, fluorescence microscopy and quantitative PCR analyses showed up to 25-fold and 10-fold higher expression levels with VAST compared to AGROBEST and Fast-TrACC, respectively (**Figure 2C**), which is reproducible using different *Agrobacterium* strains (GV3101 and C58C1, **Figure 3**). Additionally, VAST does not cause severe plant stress or tissue damage (**Supplementary Figure 7**), as indicated by low expression of the canonical stress markers ADH (**Figure 2D**), PR1 (**Supplementary** Figure 3), and CAT (**Supplementary Figure 1**5D) and low reactive oxygen species accumulation (**Supplementary** Figure 8**, X**), preserving overall plant health and enabling longer-term studies up to 16 dpi (**Supplementary Figure 9,10**). We have also shown that VAST can be applied to both plate-and soil-grown plants (**Supplementary Figure 5**). Detailed analysis of reporter expression at the single-cell resolution reveal transformation efficiencies up to 90% in some Arabidopsis seedlings with an average of 60% in all transformed seedlings, and up to 40% in Setaria with an average of 23% (**Supplementary Figure 11,15**).

The core technological advance of our VAST is the optimization of plant pre-treatment with vacuum and sonication prior to *Agrobacterium* co-cultivation. This increases bacterial access to internal tissues, i) improving transient expression efficiency in plants like Arabidopsis that have been previously transiently transformed, and ii) enabling robust transient transformation in previously recalcitrant species, including monocots. VAST is synergistic with existing approaches that enhance transformation efficiency through *Agrobacterium* strain improvement or modified *Agrobacterium* growth and/or plant handling protocols. Also, any application demonstrated in prior transient *Agrobacterium* assays should remain feasible with VAST, with the added benefit of higher expression efficiency.

Importantly, VAST overcomes some of the limitations associated with protoplast-based transient systems. Despite their widespread use, protoplasts cannot be obtained from all plant species, and they suffer from limited cell viability and timespan, loss of cell type identity, and absence of systemic biological interactions. In contrast, VAST enables sustained expression in intact seedlings over extended periods (**Supplementary Figure 9,10**), facilitating physiological and developmental studies that are biologically relevant and previously unfeasible using protoplasts, such as in our case study of nitrate availability response. Our demonstration of the nitrate-response assay highlights the functional utility and precision of VAST, showcasing its capability to interrogate plant responses to environmental signals under relevant conditions (**Figure 6**). This method allowed for the accurate detection of nitrate-specific responses in *A*. *thaliana* and *S*. *italica* across 9 days and revealed interesting and previously unknown function conversation and divergence in these plants’ nitrate availability response (**Figure 6** and **Supplementary Figure 16**).

Despite the promising advantages, VAST faces certain limitations, notably species-specific optimization requirements, as exemplified by differential responses observed among tested monocot and eudicot species. In particular, transient expression in wheat remains relatively low compared to other species, suggesting that additional optimization in *A. tumefaciens* strain and OD, sonication intensity, vacuum conditions, and co-cultivation parameters may be necessary.

Future research will explore compatibility with CRISPR/Cas-mediated genome editing or base-editing systems, where transient delivery of editing components via VAST could enable precise genomic edits without transgene integration, thus simplifying the generation of transgene-free plants. Additionally, given that the core improvements in VAST derive from optimized physical treatments facilitating enhanced *A. tumefaciens* infiltration, future studies may further boost efficiency by integrating recently developed engineered *A. tumefaciens* strains, such as the ones based on the hypervirulent GV2260 background with additional virulence genes, disabled recombination machinery, and controlled copy-number plasmids, amongst many other new developments.^16–19^ Leveraging these improved strains and protocols within the VAST framework may further boost transformation efficiencies, broadening its applicability and impact across recalcitrant plant species.

In conclusion, VAST offers a versatile, efficient, and biologically relevant alternative for transient transformation, addressing some of the critical limitations of current methods. It opens avenues for rapid functional genomic studies, synthetic biology, and potential improvements in stable transformation protocols across diverse plant species. Further optimization and integration with targeted genome engineering approaches promise to expand VAST’s utility, paving the way for transformative plant biotechnology applications.

## Methods

### DNA Constructs

Constructs were designed using SnapGene software (SnapGene, GSL Biotech). All plasmids were assembled through BsaI/SapI-mediated Golden Gate loop assembly. A comprehensive list of standard parts and assembled expression constructs is provided in **Supplementary Table 1**. Plasmids and sequence information are available through Addgene.

### Seed Sterilization and Germination

Col-0 *A. thaliana* seeds (Meyerowitz Lab, Caltech) were surface-sterilized by exposure to chlorine gas. Chlorine gas was produced by mixing 100 mL of bleach and 3 mL of 6N hydrochloric acid. The mixture was placed in a desiccator within a chemical fume hood, and the seeds were exposed to the chlorine gas for at least 3 hours. After sterilization, seeds were either stored or used immediately, maintaining sterile conditions throughout. Seeds were germinated on ½ MS plates containing 0.5x Murashige and Skoog Basal Salt Mixture (M524, PhytoTech Labs), 0.05% MES (w/v), 0.7% Phytagel (P8169, Millipore Sigma), pH 5.7, and incubated in a growth chamber at 22°C with a 16/8 h light/dark cycle for 8 days (5 days after germination). For soil-grown plant transformation, soil was prepared in a 20-liter plastic bin by mixing 6 parts Sunshine Mix (Sunshine^®^ Mix #5, sungro), 2 parts Perlite, and 2 parts Vermiculite (3:1:1 ratio by volume). The components were thoroughly mixed by hand until uniform. To fertilize, 9 g of Micromax (ICL) (∼7 mL) and 31 g of Osmocote 14-14-14 (LAWN SYNERGY) (∼35 mL) were added and mixed evenly. Prepared soil was transferred into trays and gently compacted. Sterilized *Arabidopsis thaliana* Col-0 seeds were evenly sown and grown at 22°C under a 16 h light/8 h dark cycle for 8 days. Before co-culture with *Agrobacterium tumefaciens*, seedlings were carefully removed and washed three times with sterile Milli-Q water to eliminate residual soil and microbial contaminants.

Transformation was conducted on several other plant species, including *Solanum lycopersicum* (tomato, cultivar Micro-Tom), *Brassica rapa* (accession Ames 30080), *Medicago sativa* (cultivar Bulldog 805), *Setaria italica* (foxtail millet, cultivar Manta Siberian), *Panicum virgatum* (switchgrass, cultivar Dacotah), and *Triticum aestivum* (wheat, soft white). All seeds, except for those of *T. aestivum*, were surface sterilized using 33% bleach (v/v) with 0.1% Tween-20 for 6 minutes. *T. aestivum* seeds were sterilized in 50% bleach (v/v) with 0.1% Tween-20 for 15 minutes. Following sterilization, the seeds were rinsed four to five times with sterile water. The sterilized seeds were then placed on ½ MS plates and maintained in a growth chamber set at 22 °C with a 16/8-hour light/dark cycle. *B. rapa* and *T. aestivum* seeds were incubated in the dark at 4 °C for 2 days to synchronize germination before being transferred to the growth chamber. Germination times and growth rates varied among species on ½ MS plates. Seedlings with fully expanded cotyledons and visible true leaves were used for transformation.

### Preparation of *A. tumefaciens* Cultures

*Agrobacterium-mediated* transformation using the VAST method was performed using *Agrobacterium tumefaciens* strain GV3101 (1282PS-12, Intact Genomics), carrying pSoup and the plasmid of interest, for eudicot species including *A. thaliana, S. lycopersicum*, *B. rapa*, and *M. sativa*. For *A. thaliana*, we additionally used the C58C1(pTiB6S3ΔT)^H^ strain, a kind gift from Dr. Lai. *A. tumefaciens* strain AGL1 (1283PS-12, Intact Genomics) and EHA105 (1084-06, Intact Genomics) were used to infect monocot plants, including *S. italica, P. virgatum, and T. aestivum. A. tumefaciens* cultures were inoculated from glycerol stocks into 2X-YT liquid medium (22712-020, Thermo Fisher Scientific) containing the appropriate antibiotics and incubated at 28°C with shaking at 200 rpm for 16-20 hrs. The cultures were then harvested by centrifugation at 8,000g for 5 minutes, and the resulting pellet was resuspended to an OD_600_ of 0.3 in AB-MES solution (17.2 mM K_2_HPO_4_, 8.3 mM NaH_2_PO_4_, 18.7 mM NH_4_Cl, 2 mM KCl, 1.25 mM MgSO_4_, 100 μM CaCl_2_, 10 μM FeSO_4_, 50 mM MES, 2% glucose (w/v), 200 μM acetosyringone, pH 5.5) (Wu et al., 2014), without antibiotics, and incubated overnight at 28 °C with shaking at 200 rpm. The following day, co-culture media consisting of a 50:50 (v/v) mix of AB-MES solution and ½ MS liquid plant growth medium (0.5x Murashige and Skoog Basal Salt Mixture (M524, PhytoTech Labs), 0.05% MES (w/v), 0.5% sucrose (w/v), pH 5.7), supplemented with 200 μM acetosyringone (40100297-2, PlantMedia), were prepared. *A. tumefaciens* cultures were pelleted and resuspended in co-culture media at different OD_600_ (0.14 or 0.4) for infection.

### Co-cultivation of *A. tumefaciens* and Seedlings

Five-day-old *A. thaliana* seedlings, at the stage where two small true leaves are emerging, were immersed in *A. tumefaciens* suspension in co-culture media (OD_600_ = 0.14) in 1.5 mL microcentrifuge tubes, with 10–15 seedlings per tube. The tubes were sonicated for 20 seconds using an ultrasonic cleaner (CPX-952-116R, Branson Bransonic) to enhance *A. tumefaciens* infiltration. Seedlings were then transferred to six-well plates (FB012917, Fisher Scientific), and an additional 1 mL of *A. tumefaciens* suspension was added to each well, bringing the total volume to 2 mL.

The plates were placed in a vacuum chamber (24988-233, VWR), and three vacuum cycles were performed. Each cycle involved maintaining the plates under vacuum for 5 minutes, followed by rapid pressure release. After each cycle, the plates were briefly swirled to ensure even distribution of the *A. tumefaciens* suspension. Following vacuum infiltration, the seedlings were returned to the growth chamber and co-cultivated with *A. tumefaciens* for 2 days.

Following co-cultivation, seedlings were transferred to 100 mm Petri dishes (08-757-100D, Fisher Scientific) containing sterile water to remove residual *A. tumefaciens*. After washing, the seedlings were transferred to ½ MS plates supplemented with 100 μM timentin (NC0091860, Fisher Scientific) to eliminate remaining *A. tumefaciens* and maintain sterility for subsequent analyses. Seedlings were returned to the growth chamber, and the transformation step was completed.

#### AGROBEST

The optimized method developed by Wu et al., known as AGROBEST^4^, was performed as follows. *A. tumefaciens* was freshly streaked from a -80°C glycerol stock onto an agar plate and incubated at 28°C for 2 days. A single fresh colony from the plate was used to inoculate 5 mL of 523 liquid medium containing the appropriate antibiotics. The culture was grown at 28°C with shaking at 200 rpm for 20–24 hours. To induce *A. tumefaciens* vir gene expression, cells were pelleted by centrifugation and resuspended to an OD_600_ of 0.2 in AB-MES containing 200 μM of acetosyringone. The cultures were incubated at 28°C with shaking at 200 rpm for 12–16 hours. After this step, the cultures were centrifuged again and resuspended to an OD_600_ of 0.02 in a 50:50 (v/v) mixture of AB-MES solution and ½ MS liquid plant growth medium. Seedlings were geminated and maintained in a growth chamber at 22°C under a 16/8-hour light/dark cycle. After germination, the seedlings were grown for an additional five days in ½ MS medium (eight days in total). Subsequently, the ½ MS medium was removed, and the prepared *A. tumefaciens* culture was added to the wells for co-culture. The seedlings were incubated with *A. tumefaciens* for two days, after which they were thoroughly washed with sterile water to remove residual bacteria. Following the washing step, the seedlings were transferred to 6-well plates containing 1 mL of fresh ½ MS liquid medium per well (5 seedlings per well) supplemented with 100 μM timentin to eliminate any remaining *A. tumefaciens*.

### Fast-TrACC

The optimized method developed by Nasti et al., known as Fast-TrACC^6^, was performed as follows. *A. tumefaciens* cultures (strain GV3101) were prepared by initially growing them overnight (8–12 hours) in Luria broth supplemented with the appropriate antibiotics at 28°C. The cultures were then harvested by centrifugation and resuspended in AB-MES solution containing 200 μM acetosyringone to an OD_600_ of 0.3, followed by an additional overnight incubation. After this step, the cultures were centrifuged again and resuspended to an OD_600_ of 0.14 in a 50:50 (v/v) mixture of AB-MES solution and ½ MS liquid plant growth medium.

*A. thaliana* seeds were sterilized by washing with 70% ethanol for 1 minute, followed by treatment with 50% (v/v) bleach for 5 minutes, and then rinsed five times with sterile water. Sterilized seeds were transferred to 6-well plates, with approximately five seeds per well in 2 mL of ½ MS liquid medium and maintained in a growth chamber at 22°C under a 16/8-hour light/dark cycle. After germination, the seedlings were grown for an additional five days in ½ MS medium (eight days in total). Subsequently, the ½ MS medium was removed, and the prepared *A. tumefaciens* culture was added to the wells for co-culture. The seedlings were incubated with *A. tumefaciens* for two days, after which they were thoroughly washed with sterile water to remove residual bacteria. Following the washing step, the seedlings were transferred to 6-well plates containing 2 mL of fresh ½ MS liquid medium per well (5 seedlings per well) supplemented with 100 μM timentin to eliminate any remaining *A. tumefaciens*.

### Co-cultivation of *A. tumefaciens* with seedlings of other species

The VAST transient transformation method was also tested in several other species, including three eudicot species (*S. lycopersicum*, *B. rapa*, and *M. sativa*) and three monocot species (*S. italica*, *P. virgatum*, and *T. aestivum*), with some modifications in the corresponding species which are described below and summarized in Supplementary Table 3.

### Co-cultivation of *A. tumefaciens* and *S. lycopersicum* seedlings

*S. lycopersicum* seedlings were transferred into 50 mL Falcon tubes and immersed in about 8 mL of *A. tumefaciens* GV3101 suspension in co-culture media (OD_600_ = 0.4), with 7-8 seedlings per tube. The tubes were sonicated for 30 seconds using an ultrasonic cleaner (CPX-952-116R, Branson Bransonic) and subsequently placed in a vacuum chamber (24988-233, VWR) for three cycles of vacuum infiltration. After each cycle, the tubes were briefly shaken. Seedlings were then transferred to six-well plates (FB012917, Fisher Scientific) containing 4 mL *A. tumefaciens* suspension per well, returned to the growth chamber, and co-cultivated with *A. tumefaciens* for 2 days.

### Co-cultivation of *A. tumefaciens* and *B. rapa* seedlings

*B. rapa* seedlings underwent a similar process but were sonicated for 15 seconds followed by three cycles of vacuum infiltration. Three cycles of vacuum infiltration only sometimes also yield high YPET expression, however, the efficiency varies a lot between seedlings (Supplementary Figure S5B). After vacuum infiltration, seedlings were transferred to six-well plates containing 4 mL *A. tumefaciens* suspension in each well, returned to the growth chamber and co-cultivated with *A. tumefaciens* for 1 day.

### Co-cultivation of *A. tumefaciens* and *M. sativa* seedlings

*M. sativa* seedlings underwent a similar process but were sonicated for 45 seconds followed by four cycles of vacuum infiltration. After vacuum infiltration, seedlings were transferred to six-well plates containing 4 mL *A. tumefaciens* suspension in each well, returned to the growth chamber and co-cultivated with *A. tumefaciens* for 2 days.

### Co-cultivation of *A. tumefaciens* and monocot seedlings

Monocot seedlings were transferred into 50 mL Falcon tubes and immersed in about 8 mL of *A. tumefaciens* AGL1 suspension in co-culture media (OD_600_ = 0.4), with 7-8 seedlings per tube. The tubes were sonicated for 30 seconds followed by four cycles of vacuum infiltration. The 45 seconds of sonication treatment followed by four cycles of vacuum infiltration has also been tested in the monocot *P. virgatum*. The increase in sonication duration leads to higher YPET expression but also results in more tissue damage (Supplementary Figure S6B). After vacuum infiltration, seedlings were transferred to six-well plates containing 4 mL *A. tumefaciens* suspension in each well, returned to the growth chamber and co-cultivated with *A. tumefaciens* for 2 days.

Following co-cultivation, all the seedlings were rinsed with sterile water to remove residual *A. tumefaciens* and then transferred to ½ MS plates supplemented with 100 μM timentin (NC0091860, Fisher Scientific) to eliminate remaining *A. tumefaciens* and potential contaminants. The seedlings were then maintained in the growth chamber for subsequent analyses.

### Imaging and Analysis

All the transformed seedlings were imaged at 4-5 or 10 dpi. YPET fluorescence, chlorophyll fluorescence, and bright-field images of seedlings were captured using a Revolve R4 microscope (Discover Echo). Seedling leaves were also imaged using a Zeiss LSM 880 upright confocal microscope or a STELLARIS 8 inverted confocal microscope. YPET was excited using a 514-nm laser line, and the emission was collected from 508-553 nm. Images were processed using Fiji/Image J.

### Transient Expression of Luciferase Constructs and Luciferase Assay

Nitrate-responsive constructs were transformed into 5-day-old *A. thaliana* seedlings. For the kinetics assay, 2 days after co-cultivation with *A. tumefaciens*, seedlings were washed and transferred to ½ MS plates (M531 without nitrogen source, PhytoTech Labs), supplemented with 10 mM KCl. The seedlings were incubated for 3 days in nitrogen free plates to allow the activity of the NRP promoter to return to basal levels. Following this, the seedlings were transferred to ½ MS plates (M531, PhytoTech Labs), supplemented with 10 mM KNO_3_, and luminescence signals were measured daily. To evaluate the sensor’s response and specificity, seedlings were washed two days after co-cultivation with *A. tumefaciens* and transferred to ½ MS plates (M531, PhytoTech Labs) with different treatments. For assessing the sensor’s response, the treatments included 0, 1, or 10 mM KNO₃, with additional KCl added as necessary to maintain a consistent potassium concentration. For evaluating sensor specificity, treatments of 10 mM KCl, glutamine, NH₄Cl, KNO₃, or NH₄NO₃ were applied. The seedlings were then incubated for three additional days, after which luminescence signals were measured.

Luminescence signals were quantified by collecting five seedlings per sample into 2 mL tubes (15-340-162, Fisher Scientific) containing 2.8 mm ceramic beads (15-340-160, Fisher Scientific). Followed by immediate flash-freezing in liquid nitrogen, frozen samples were ground using a mini-bead beater tissue homogenizer (15-340-164, Fisher Scientific) for 40 seconds at maximum speed. Subsequently, 100 µL of 1x cell lysis buffer (E1531, Promega) was added to each tube, and the samples were vigorously vortexed until fully dissolved, ensuring no clumps remained. The tubes were placed on ice for 10 minutes, then centrifuged at 5,000g for 5 minute to pellet the cell debris.

Luminescence was measured using the Nano-Glo® Luciferase Assay System (N1110, Promega), following the manufacturer’s instructions with minor modifications. A 15 μL aliquot of protein extract from the previous step was added to 30 μL of Nano-Glo reagent per well in a 96-well plate (07-200-627, Fisher Scientific). Luminescence was measured using a Tecan Spark plate reader at 25°C, with GeNL (520–590 nm) and NLuc (430–485 nm) filters, and an integration time of 1 second per well. To calculate signal deconvolution coefficients, total, GeNL, and NLuc luminescence were measured, and a Promega ChromaLuc Technology calculator spreadsheet (Promega Technical Manual TM062) was used.

### Plant RNA Extraction and Quantitative RT-PCR

RNA extraction and RT-qPCR were performed to quantify gene expression in *A. thaliana* seedlings using the following commercially available kits: the RNeasy Plant Mini Kit (74904, QIAGEN) for total RNA extraction, TURBO™ DNase (AM2238, Thermo Fisher Scientific) for removing residual DNA, M-MLV Reverse Transcriptase (28025013, Thermo Fisher Scientific) to reverse transcribe RNA into cDNA, and PowerUp SYBR Green Master Mix (A25776, Thermo Fisher Scientific) for qPCR. All primers used are listed in **Supplementary Table 2** and were ordered from IDT. The qPCR was conducted with an annealing temperature of 60°C for 40 cycles.

qPCR data were analyzed using the ΔCt method to obtain normalized gene expression relative to the *EF1α* housekeeping gene. Each experiment included 3–4 biological replicates, with each replicate consisting of five plant seedlings. qPCR was performed with two technical replicates (two reactions from the same RNA batch) and averaged to calculate the final Ct values.

### DAB Staining

Hydrogen peroxide accumulation in leaves was detected using 3,3’-diaminobenzidine (DAB) staining as described by Daudi and O’Brien (2012)^21^. A DAB staining solution was freshly prepared by dissolving 50 mg of DAB in 45 mL sterile water, adjusting the pH to 3.0 with 0.2 M HCl, and adding 25 μL of Tween 20 and 2.5 mL of 200 mM Na₂HPO₄, resulting in a final concentration of 1 mg/mL DAB and 10 mM Na₂HPO₄. Seedlings were subjected to experimental treatments and then immersed in the DAB solution in 12-well plates. Vacuum infiltration was performed using a desiccator for 5 minutes to ensure solution uptake. Samples were incubated in the dark on a shaker at 80-100 rpm for 4-5 hours. Following incubation, leaves were bleached by replacing the staining solution with a boiling bleach solution (ethanol:acetic acid:glycerol, 3:1:1) for 15 minutes, then further cleared in fresh bleach solution for 30 minutes.

## Data Availability

The authors declare that the data supporting the findings of this study are available within the paper and its supplementary information files. Supplementary Table 1 contains Addgene IDs for all deposited plasmids. Source Data are provided with this paper.

## Funding

This research was supported by grants from the Henry Luce Foundation, Shurl Kay Curci Foundation, NASA Translational Research Institute for Space Health (TRISH) through Caltech Space-Health Innovation Fund, and Caltech internal seed and startup funds. EL is supported by the National Science Foundation (NSF) Graduate Research Fellowships Program (GRFP). YW is supported by the Foundation for Food & Agriculture Research (FFAR) Fellows Program and the Caltech Resnick Sustainability Institute (RSI).

## Author contributions

E.L., Y.G., and G.S.D. conceived the project, designed the study, and wrote the manuscript. E.L. and Y.G. conducted most of the experiments and performed all data analysis. Y.W. captured some of the confocal images of monocots and eudicots. T.R.K. optimized and carried out transformation for some of the monocots and eudicots, while J.J. performed some of the eudicot transformations. Y.W., J.J., and T.R.K. contributed to the functional analysis of plant nitrate response. All authors reviewed, edited, and approved the final manuscript.

## Acknowledgements

The authors gratefully acknowledge the Meyerowitz Lab at Caltech for providing *A. thaliana* Col-0 seeds and access to the Revolve R4 microscope. We also thank the Caltech Biological Imaging Facility for their expertise and support, as well as for providing access to the Zeiss LSM 880 upright confocal microscope and the STELLARIS 8 inverted confocal microscope. We further thank Dr. Lai for generously providing the *Agrobacterium tumefaciens* strain C58C1(pTiB6S3ΔT)^H^ used in our experiments.

## Declaration of interests

The authors declare that they have no known competing financial interests or personal relationships that could have appeared to influence the work reported in this paper.

## Notes

### Competing Interest Statement

The authors have declared no competing interest.

### Summary of Updates

Briefly, the revised manuscript has the below major additional data: 1. Showing VAST transformation with other Agrobacteria strains of C58C1 and EHA105 2. Systematic optimization of vacuum and sonication duration for VAST 3. Detailed efficiency analysis at spatial and temporal scales, reporting % transformation 4. Detailed damage analysis via histology, ROS measurements, more imaging, and additional qPCR stress marker gene expression levels 5. Expression of an intron-containing reporter gene with VAST 6. Showing VAST efficacy in soil-grown plants

