## Supplementary Info for "Vacuum and sonication treatment enables efficient transient gene expression in various monocot and eudicot plant seedlings"

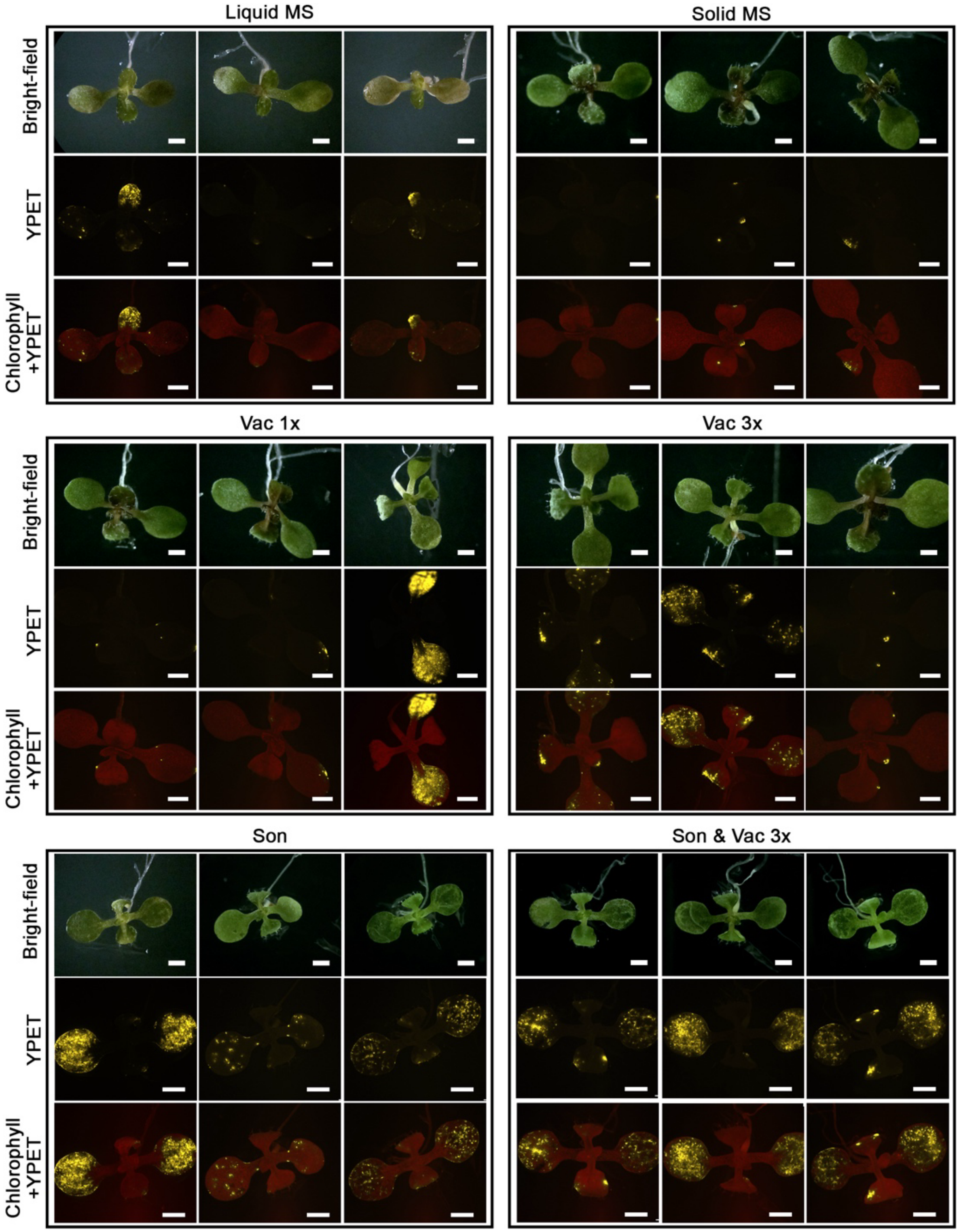


**Supplementary Figure 1. Fluorescence microscopy images of *Arabidopsis thaliana* seedlings after different treatments.** Vac 1x: Vacuum once; Vac 3x: Vacuum 3 times; Son: Sonication 20s; Son & Vac 3x: combination of sonication 20s and vacuum 3 times. Channels: Bright-field, YPET (yellow), and Chlorophyll (red) overlaid with YPET. All scale bars: 1 mm.

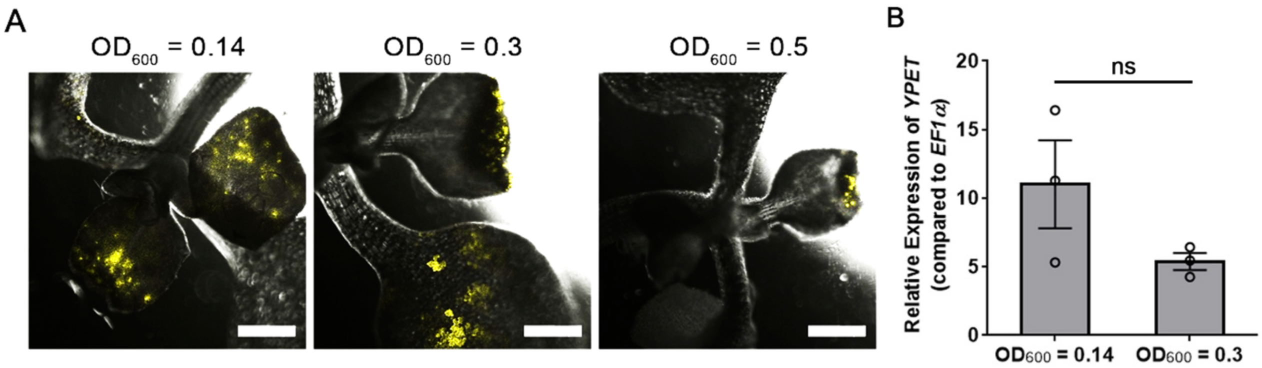


**Supplementary Figure 2. Effect of *A. tumefaciens* concentration on transient expression efficiency in *Arabidopsis thaliana*. (A)** Representative 4-dpi fluorescence microscopy images of *A. thaliana* seedlings infected with *A. tumefaciens* concentration of $OD_{600}$ = 0.14, 0.3, or 0.5. All scale bars: 500 μm. **(B)** Relative expression of *YPET* via qPCR in 4-dpi seedlings infected with *A. tumefaciens* at $OD_{600}$ = 0.14 or 0.3. N = 3 biological replicates (each with 5 seedlings). The experiment was independently repeated three times with similar results. Error bars denote standard error of mean. Statistical analysis was performed using a student's *t*-test. ns, not significant with p-value 0.1605.

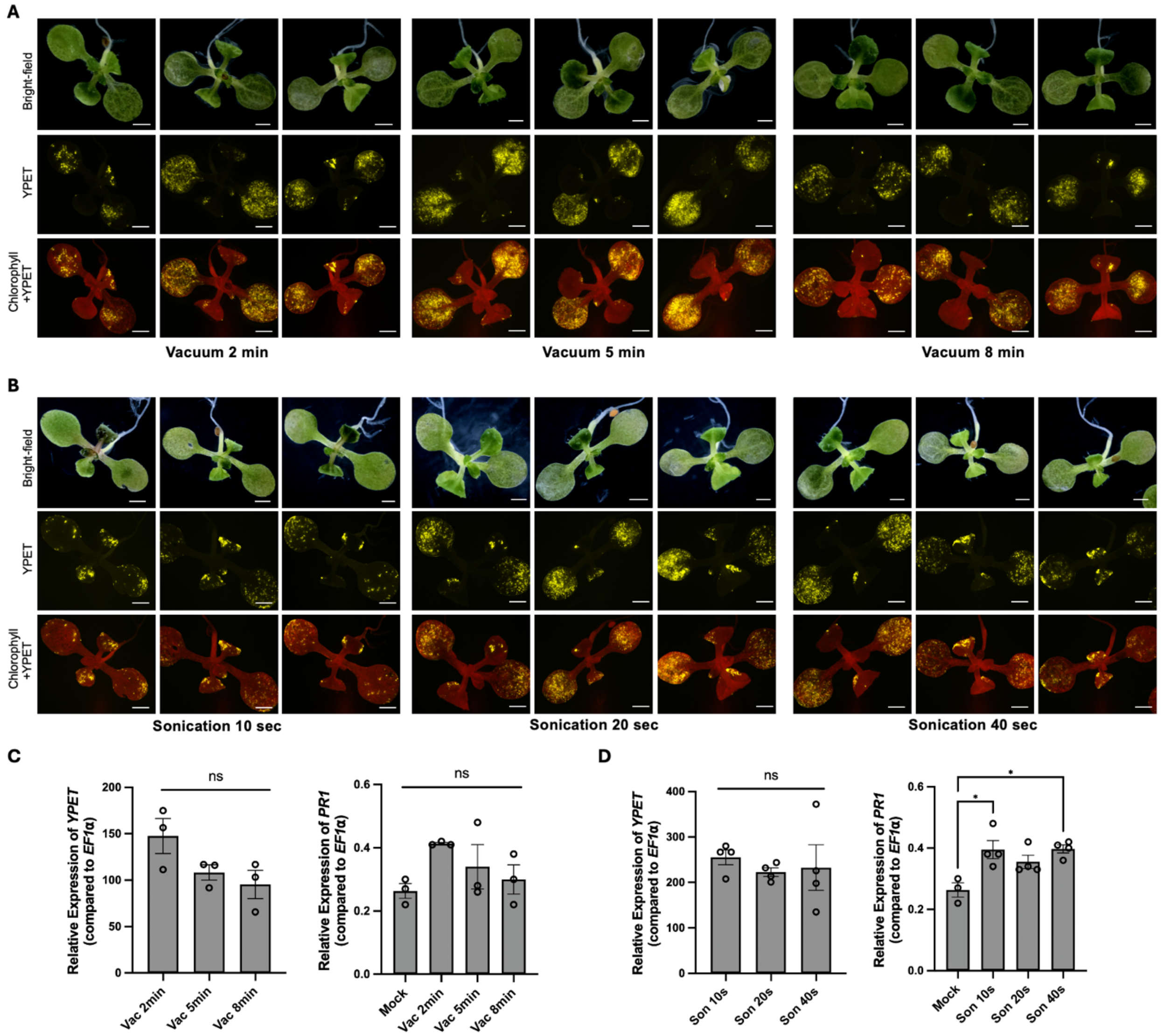


**Supplementary Figure 3. Optimization of vacuum and sonication parameters for VAST-mediated transformation efficiency and stress response in Arabidopsis. (A, B)** Representative confocal images showing YPET fluorescence in Arabidopsis seedlings subjected to different vacuum infiltration durations (2, 5, and 8 minutes) **(A)** and sonication times (10, 20, and 40 seconds) **(B)**. Top to bottom rows show bright-field, YPET fluorescence, and merged YPET + chlorophyll channels. Scale bars = 1 mm. **(C, D)** qPCR analysis of YPET (left) and PR1 (right) gene expression following vacuum **(C)** and sonication **(D)** treatments. Data are normalized to the housekeeping gene EF1α. Bars represent mean ± standard error from 3 biological replicates (each with 5 seedlings) from a representative experiment. The experiment was independently repeated three times with similar results. Statistical outliers were identified and excluded using Grubbs's test. One-way ANOVA followed by Tukey's test was used for statistical analysis. Significance levels: *$p < 0.0332$; ns, not significant.

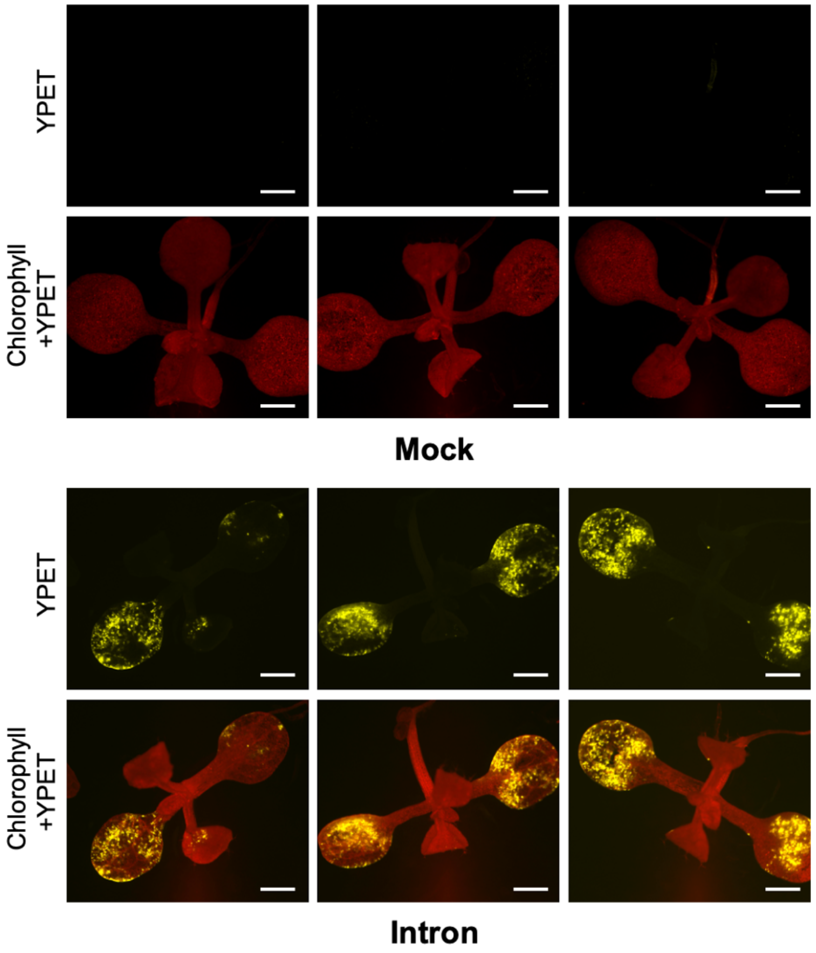


**Supplementary Figure 4. Validation of plant-specific expression of YPET**. Arabidopsis seedlings were transformed with a YPET construct containing the second intron of the ST-LS1 gene from potato to ensure expression originates from plant cells. Strong YPET fluorescence was observed in the Intron group but not in the mock-treated control, confirming plant-specific expression. Images show YPET fluorescence (top) and merged YPET + chlorophyll (bottom). Scale bars = 1 mm.

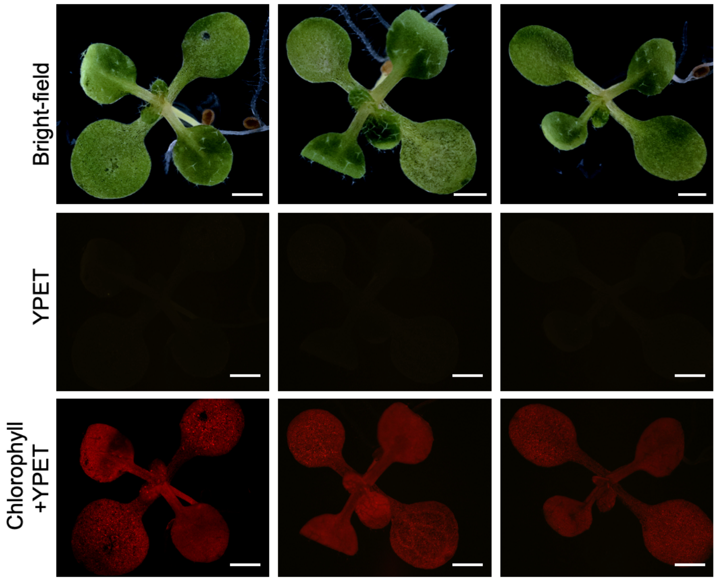


**Mock for soil-grown plants**

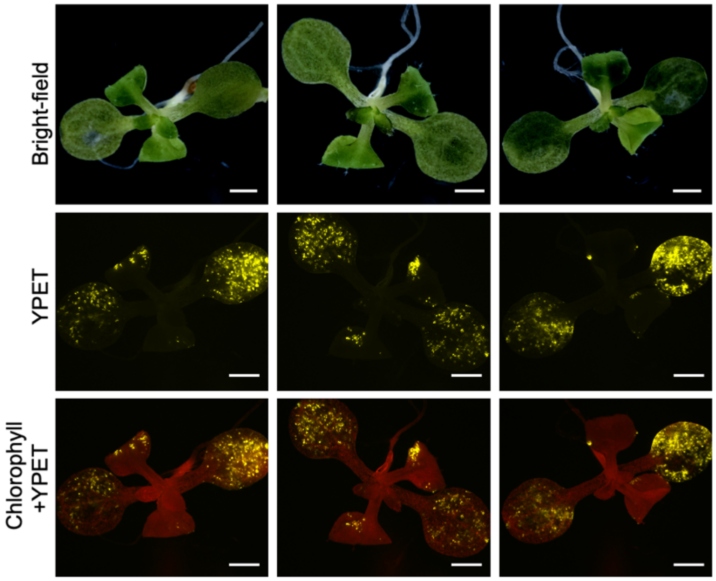


**YPET expression in soil-grown plants**

**Supplementary Figure 5. VAST enables transformation in soil-grown Arabidopsis plants.** YPET fluorescence was observed in soil-grown plants following VAST treatment, while no signal was detected in mock-treated controls. Images show bright-field (top), YPET fluorescence (middle), and merged YPET + chlorophyll (bottom). Scale bars = 1 mm.

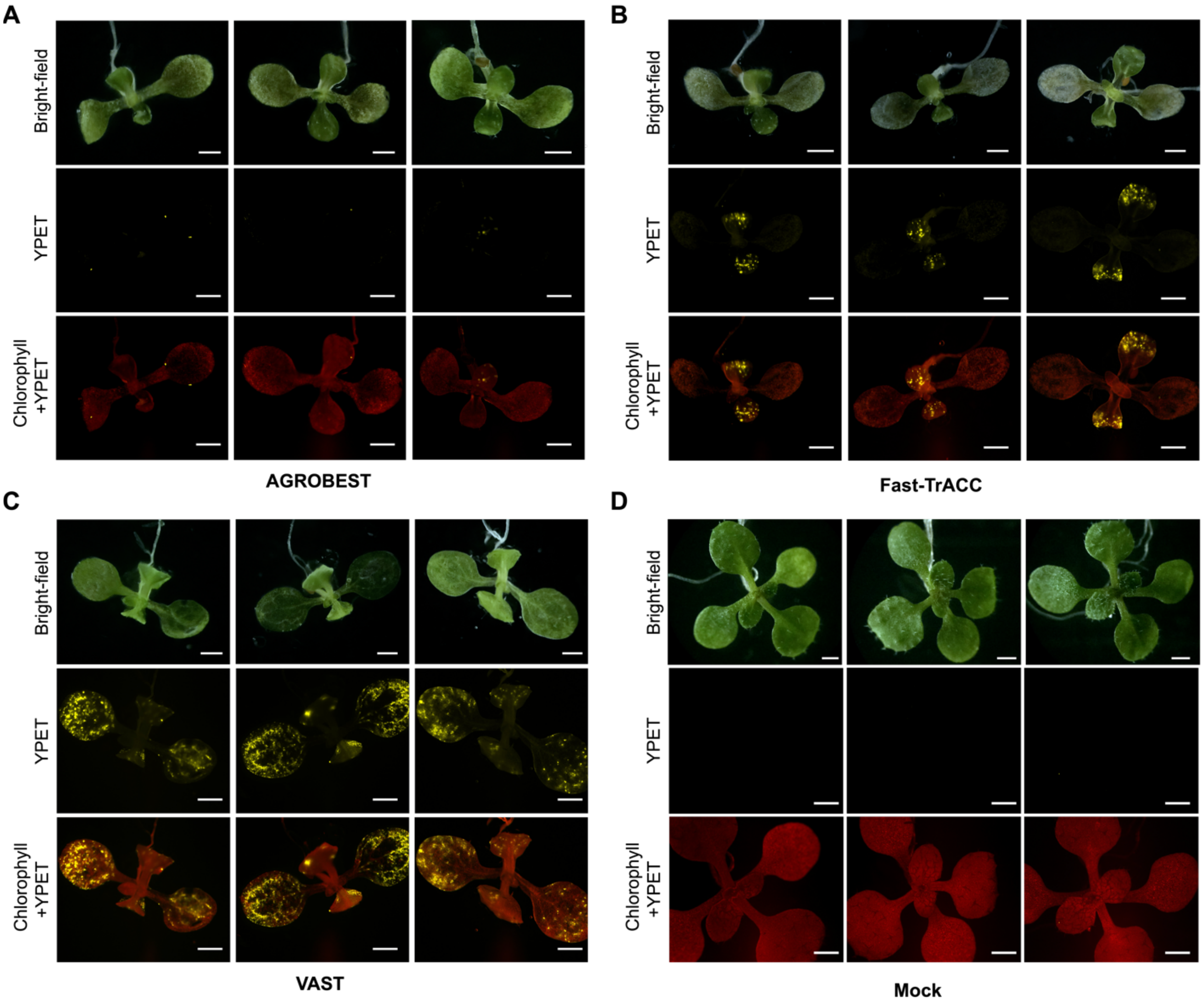


**Supplementary Figure 6.** Representative fluorescence microscopy image sets of *A. thaliana* seedlings at 4dpi using **(A)** AGROBEST, **(B)** Fast-TrACC, **(C)** VAST, and **(D)** mock treatment without *A. tumefaciens* infection (negative control). All scale bars: 1 mm. Channels: Bright-field, YPET fluorescence (yellow), and Chlorophyll (red) overlaid with YPET.

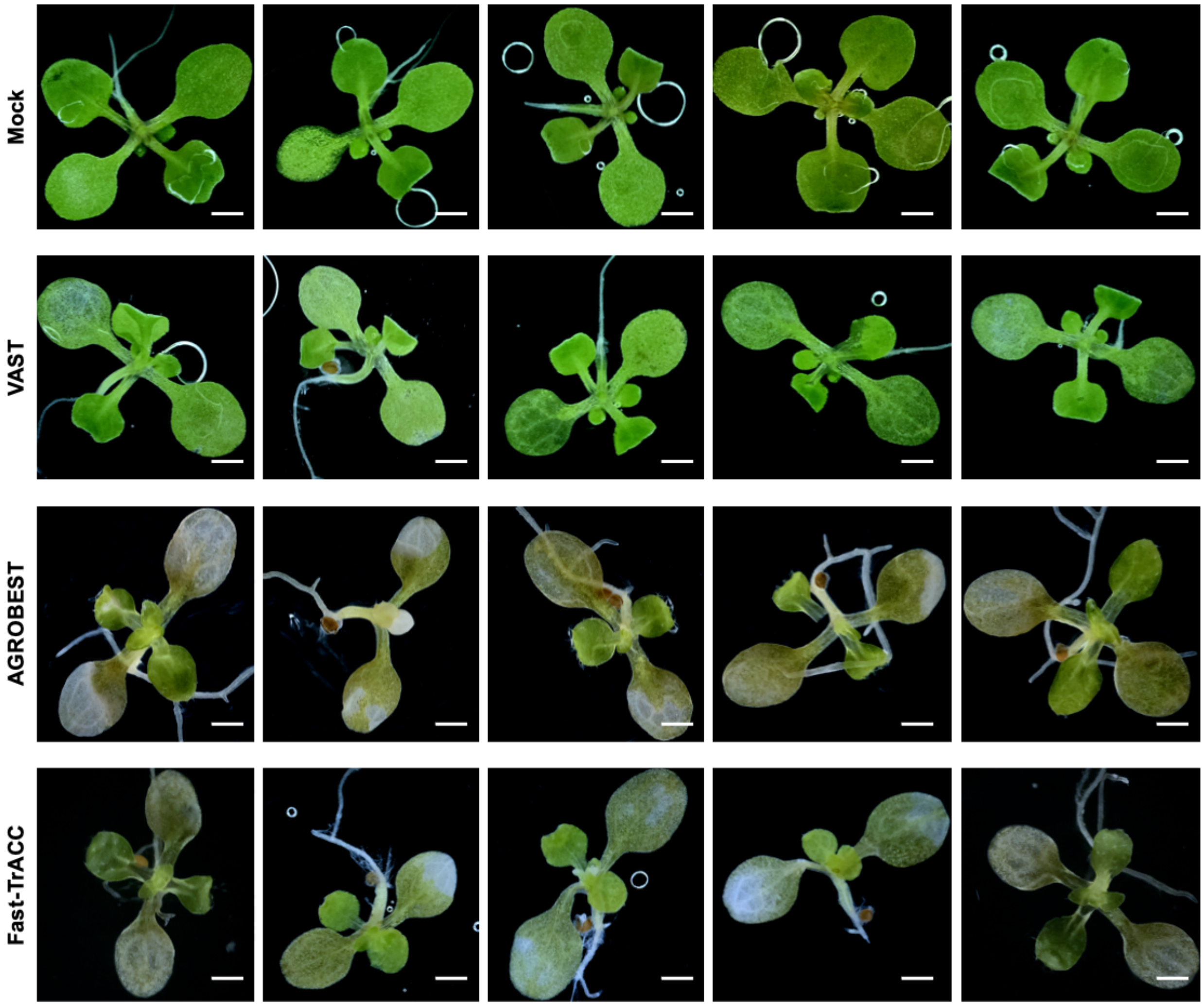


**Supplementary Figure 7.** Bright-field images showing leaf tissue condition following transformation with different methods of Mock, VAST, AGROBEST, and Fast-TrACC. Five representative seedlings are shown for each condition. Scale bars = 1 mm.

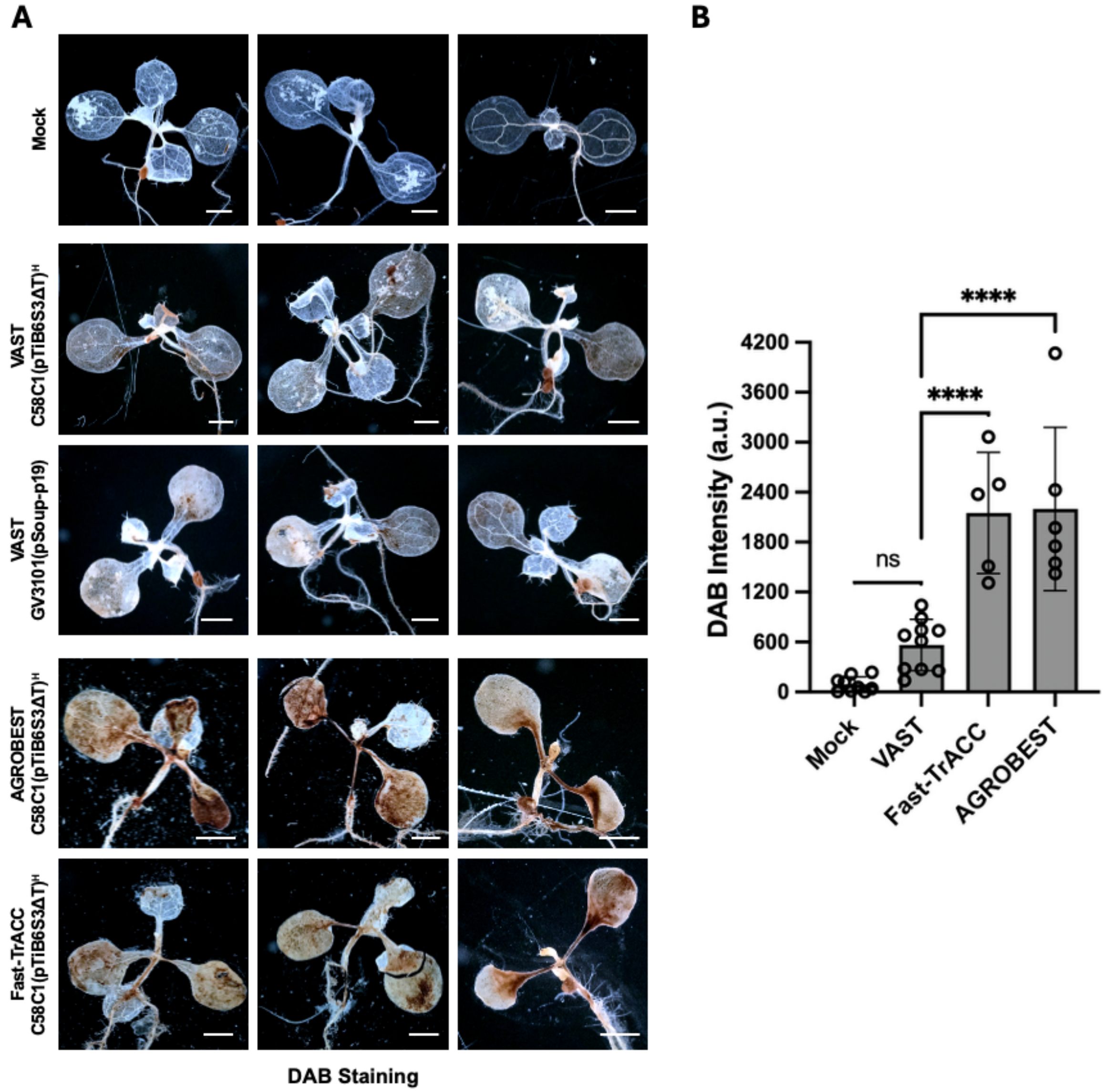


**Supplementary Figure 8. Assessment of oxidative stress levels in transformed seedlings via DAB staining. (A)** Representative images of Arabidopsis seedlings stained with DAB (3,3′-diaminobenzidine) to visualize hydrogen peroxide ($H_2O_2$) accumulation as an indicator of oxidative stress after transformation with Mock, VAST, AGROBEST, and Fast-TrACC. Brown precipitate indicates localized $H_2O_2$ accumulation. Scale bars = 1 mm. **(B)** Quantification of DAB staining intensity. Bars represent mean ± standard deviation from at least 6 biological replicates. Statistical analysis was performed using one-way ANOVA with Tukey's post hoc test. Significance levels: ****$p < 0.0001$; ns = not significant.

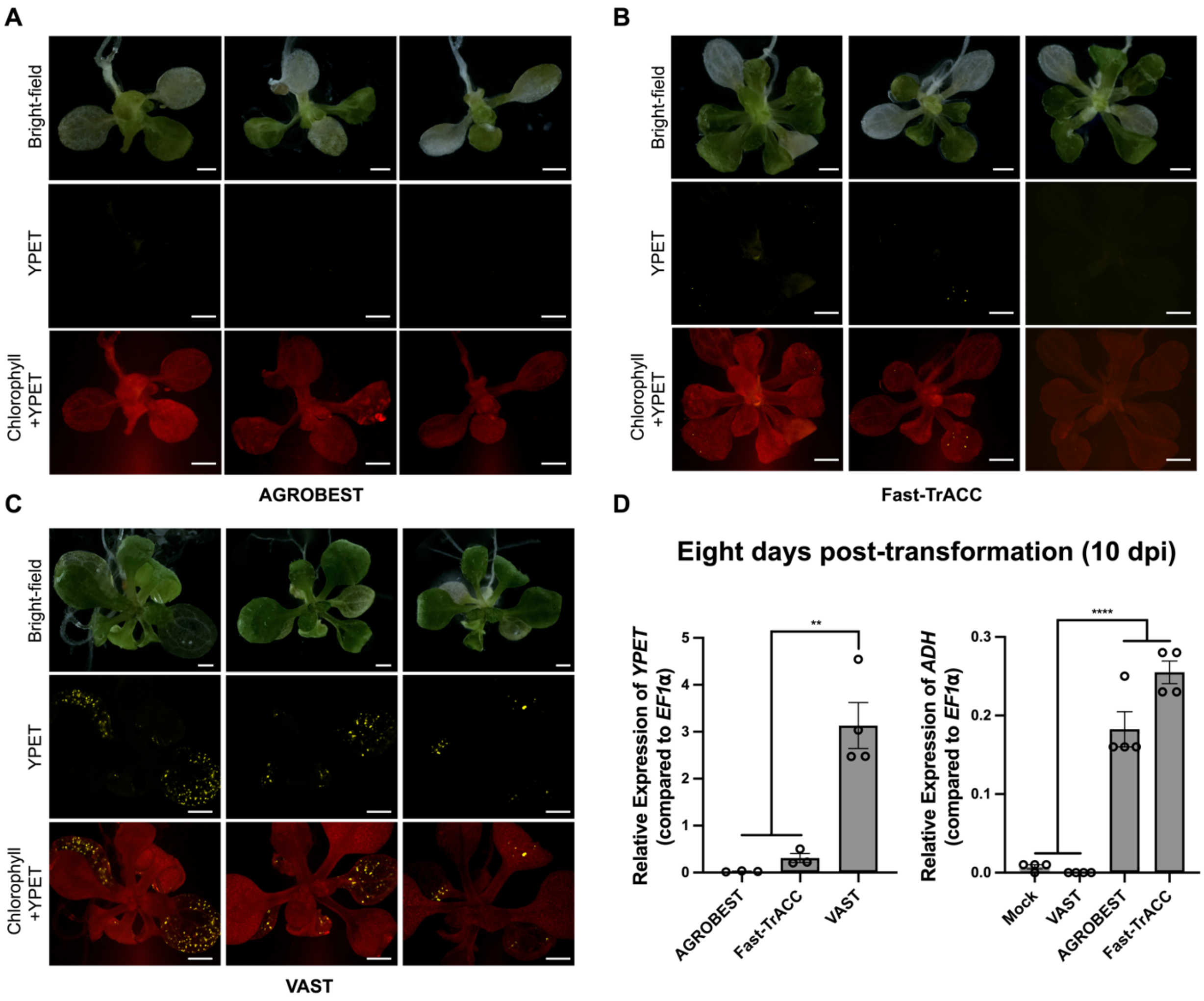


**Supplementary Figure 9.** Representative fluorescence microscopy image sets of *A. thaliana* seedlings at 10-dpi using **(A)** AGROBEST, **(B)** Fast-TrACC, and **(C)** VAST. All scale bars: 1 mm. Channels: Bright-field, YPET fluorescence (yellow), and Chlorophyll (red) overlaid with YPET. **(D)** qPCR analysis of *YPET* and *ADH* expression at 10-dpi for AGROBEST, Fast-TrACC, and VAST, normalized to the housekeeping gene *EF1α*. Data represent mean ± standard error from 4 biological replicates with 5 seedlings per replicate. The experiment was independently repeated three times with similar results. Statistical outliers were identified and excluded using Grubbs' test. One-way ANOVA with Tukey's test was used for analysis, with significance indicated as ****p $< 0.0001$, ***p $< 0.0002$, **p $< 0.0021$, *p $< 0.0332$.

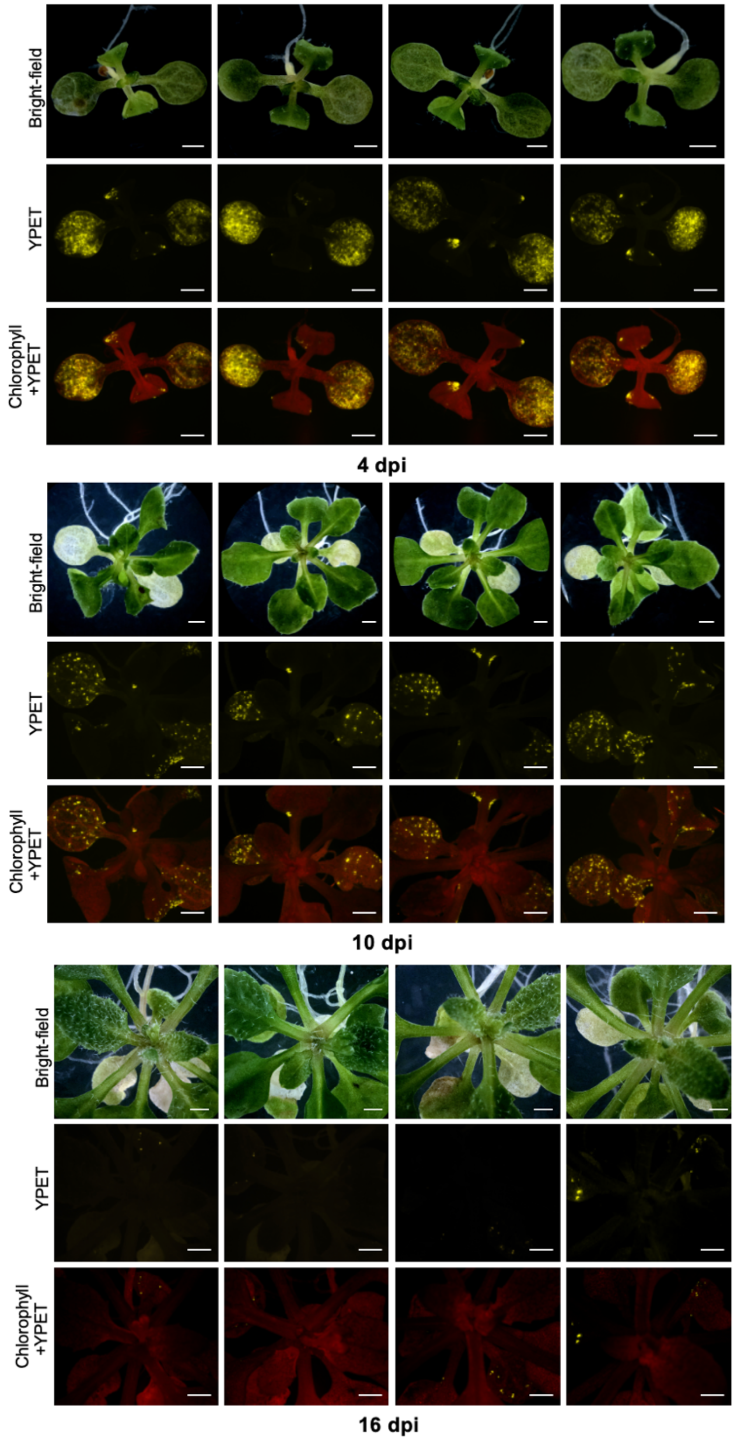


**Supplementary Figure 10. Time-course of VAST-mediated YPET expression in *Arabidopsis thaliana*.** Strong, widespread fluorescence is observed at 4 dpi, remains detectable but sparser at 10 dpi, and becomes faint by 16 dpi due to leaf expansion and emergence of new tissue. Images show bright-field, YPET, and merged YPET + chlorophyll channels. Scale bars = 1 mm.

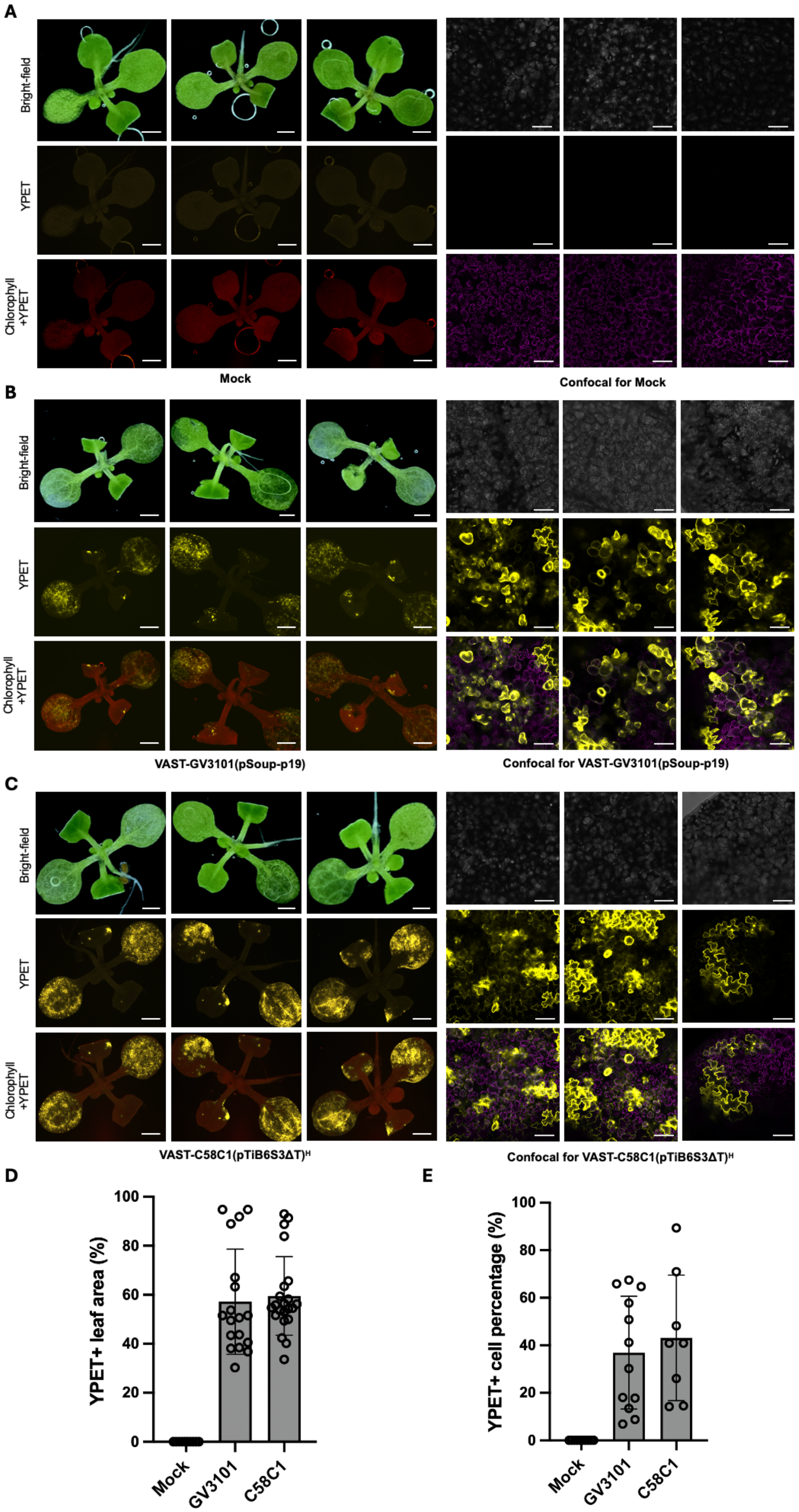
A
Bright-field
YPET
Chlorophyll +YPET
Mock
Confocal for Mock
B
VAST-GV3101(pSoup-p19)
Confocal for VAST-GV3101(pSoup-p19)
C
VAST-C58C1(pTiB6S3ΔT)H
Confocal for VAST-C58C1(pTiB6S3ΔT)H
D
YPET+ leaf area (%)
0
20
40
60
80
100
Mock
GV3101
C58C1
E
YPET+ cell percentage (%)
0
20
40
60
80
100
Mock
GV3101
C58C1

**Supplementary Figure 11. Spatial characterization and single-cell quantification of VAST-mediated expression in *Arabidopsis thaliana*. (A-C)** Representative bright-field, YPET fluorescence, and merged chlorophyll + YPET images (left) and confocal microscopy images (right) of *A. thaliana* leaves transformed with mock, VAST-GV3101(pSoup-p19), or VAST-C58C1(pTiB6S3ΔT)$^{H}$. **(D)** Quantification of YPET-positive leaf area (%), and **(E)** YPET-positive cells as a percentage of total cells, quantified from confocal images. Data represent mean ± standard deviation from at least 8 biological replicates (individual seedlings). Scale bars = 1 mm (widefield) and 100 µm (confocal).

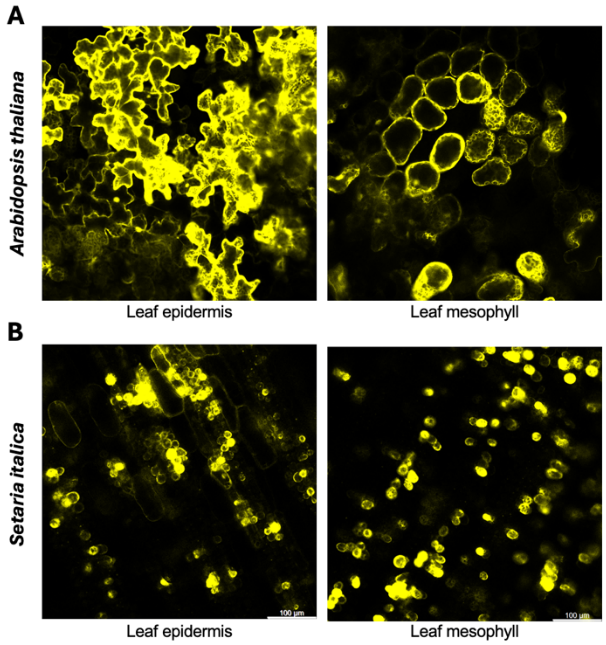


**Supplementary Figure 12.** Representative confocal microscopy images showing YPET expression in the epidermal and mesophyll layers of transformed *Arabidopsis thaliana* **(A)** and *Setaria italica* **(B)** leaves. Scale bars = 100 µm.

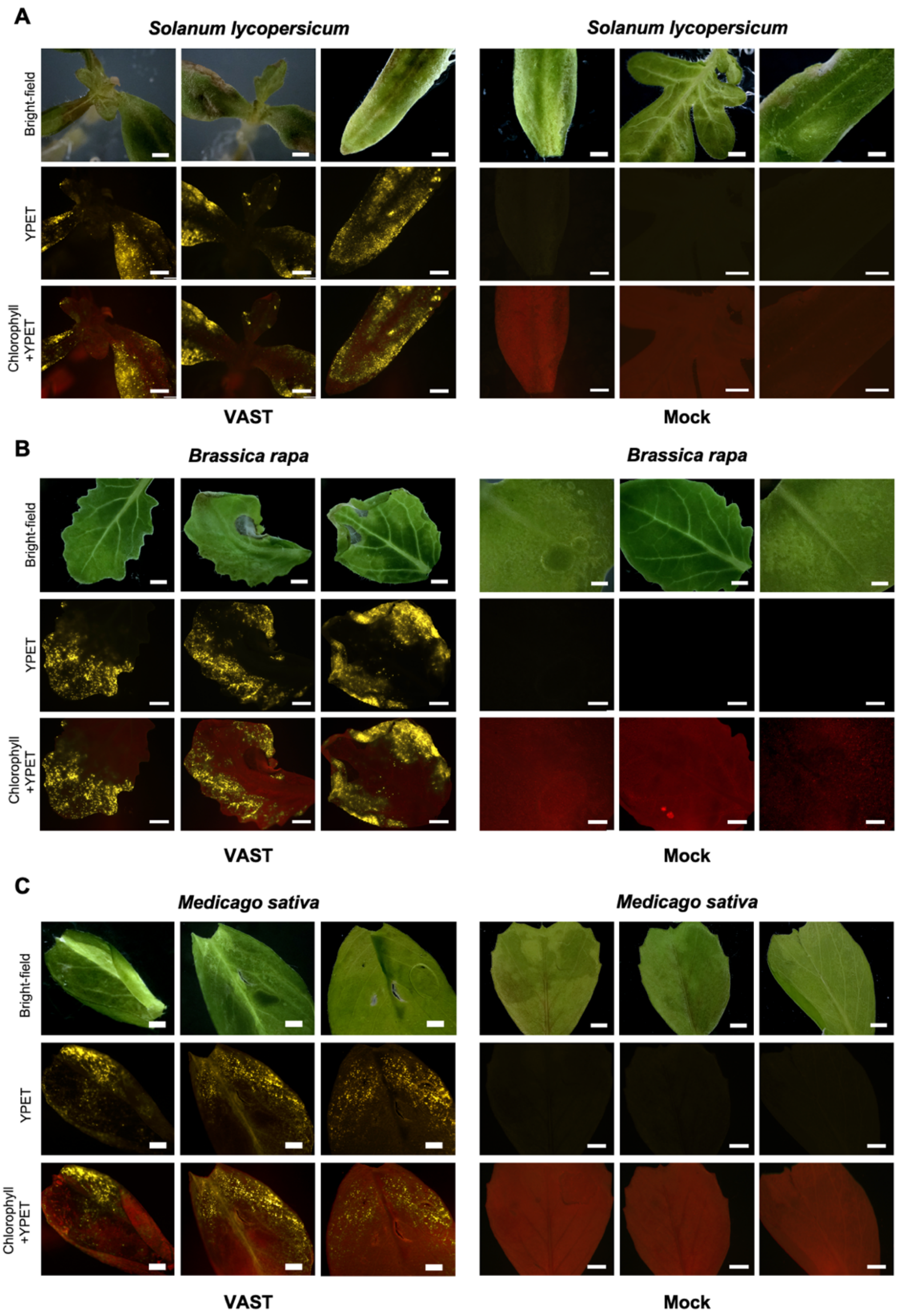


**Supplementary Figure 13.** Representative fluorescence microscopy image sets of **(A)** *Solanum lycopersicum*, **(B)** *Brassica rapa*, and **(C)** *Medicago sativa* seedlings at 4-5 dpi. The leftmost sample in (B) was treated with 3x vacuum (lacking sonication), whereas all other samples were transformed using the optimized VAST method (see **Supplementary Table 2**). All scale bars: 1 mm. Channels: Bright-field, YPET fluorescence (yellow), and Chlorophyll (red) overlaid with YPET.

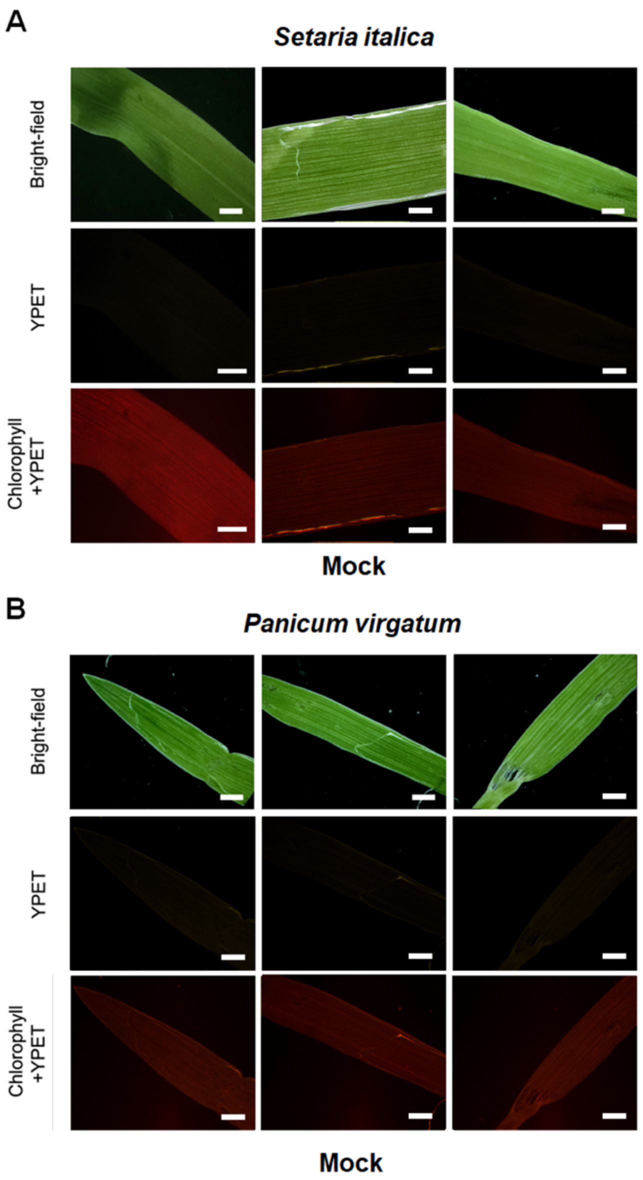


**Supplementary Figure 14.** Mock fluorescence microscopy image sets of **(A)** *Setaria italica*, and **(B)** *Panicum virgatum* at 4-dpi. All scale bars: 1 mm. Channels: Bright-field, YPET fluorescence (yellow), and Chlorophyll (red) overlaid with YPET.

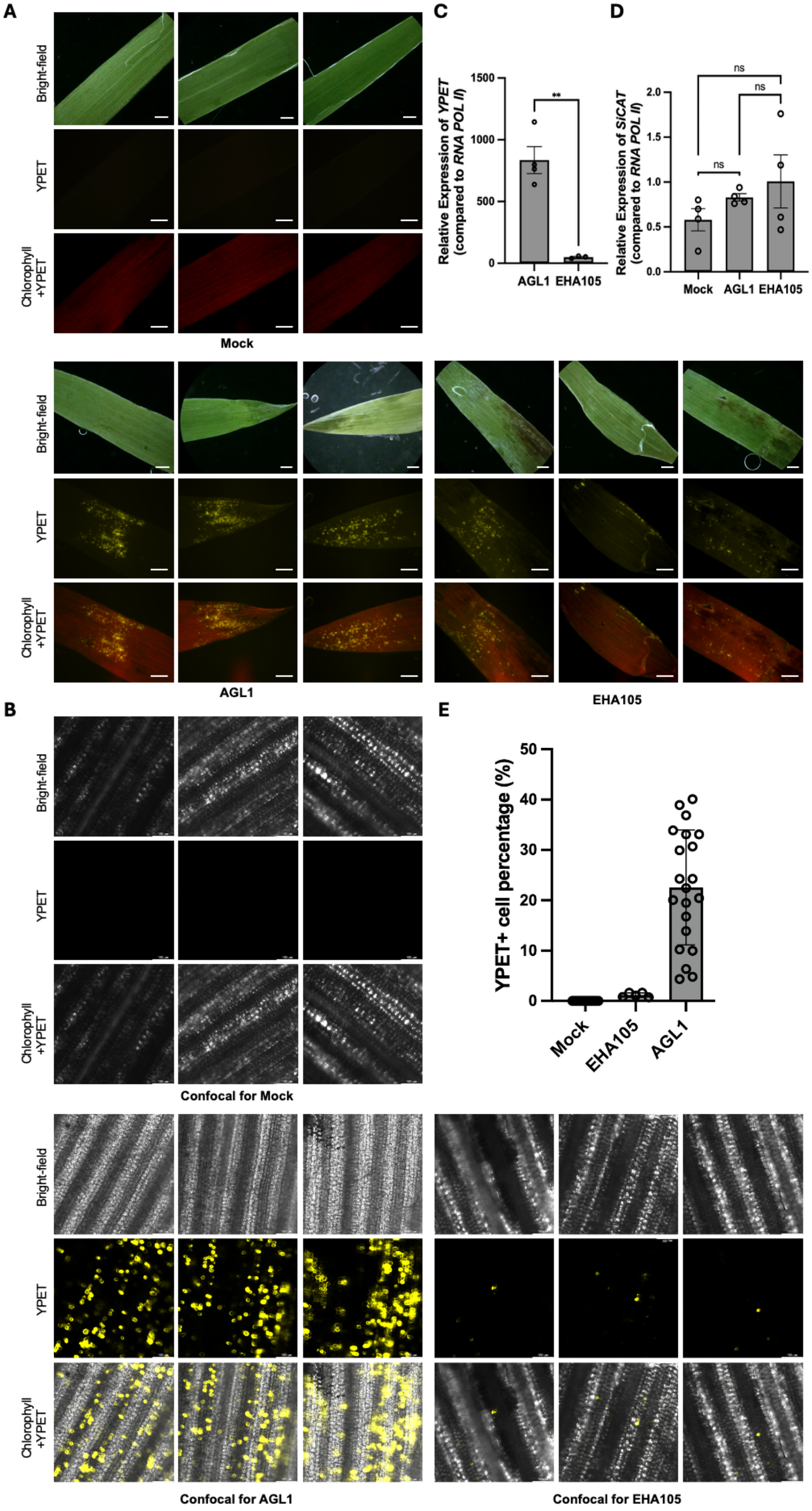
A
Bright-field
YPET
Chlorophyll +YPET
Mock
AGL1
EHA105
B
Confocal for Mock
Confocal for AGL1
Confocal for EHA105
C
Relative Expression of YPET (compared to RNA POL II)
1500
1000
500
0
**
AGL1
EHA105
D
Relative Expression of SlCAT (compared to RNA POL II)
2.0
1.5
1.0
0.5
0.0
ns
ns
ns
Mock
AGL1
EHA105
E
YPET+ cell percentage (%)
50
40
30
20
10
0
Mock
EHA105
AGL1

**Supplementary Figure 15. Comparison of *Agrobacterium tumefaciens* strains AGL1 and EHA105 for VAST-mediated transformation in *Setaria italica*. (A)** Representative bright-field, YPET fluorescence, and merged YPET + chlorophyll images of *S. italica* leaves infiltrated with mock, AGL1, or EHA105. **(B)** Representative confocal microscopy images of epidermal and mesophyll layers in AGL1-transformed leaves compared to EHA105 and mock controls. **(C)** qPCR analysis of YPET expression at 4 dpi in AGL1-treated samples compared to EHA105. **(D)** qPCR analysis of the oxidative stress marker gene *catalase* in Mock, AGL1, and EHA105-treated samples. For qPCR **(C–D)**, gene expression is compared to the housekeeping gene RNA POL II and data represent mean ± standard error from 4 biological replicates (each with 5 pooled seedlings). **(E)** Quantification of % YPET-positive cells from confocal images in 22 biological replicates (individual seedlings). Data represent mean ± standard deviation. Each dot represents one seedling. Statistical analysis was performed using one-way ANOVA with Tukey's post hoc test. Significance levels: *$p < 0.0021$; $p < 0.0332$; *ns, not significant*. Scale bars = 1 mm (widefield) and 100 μm (confocal).

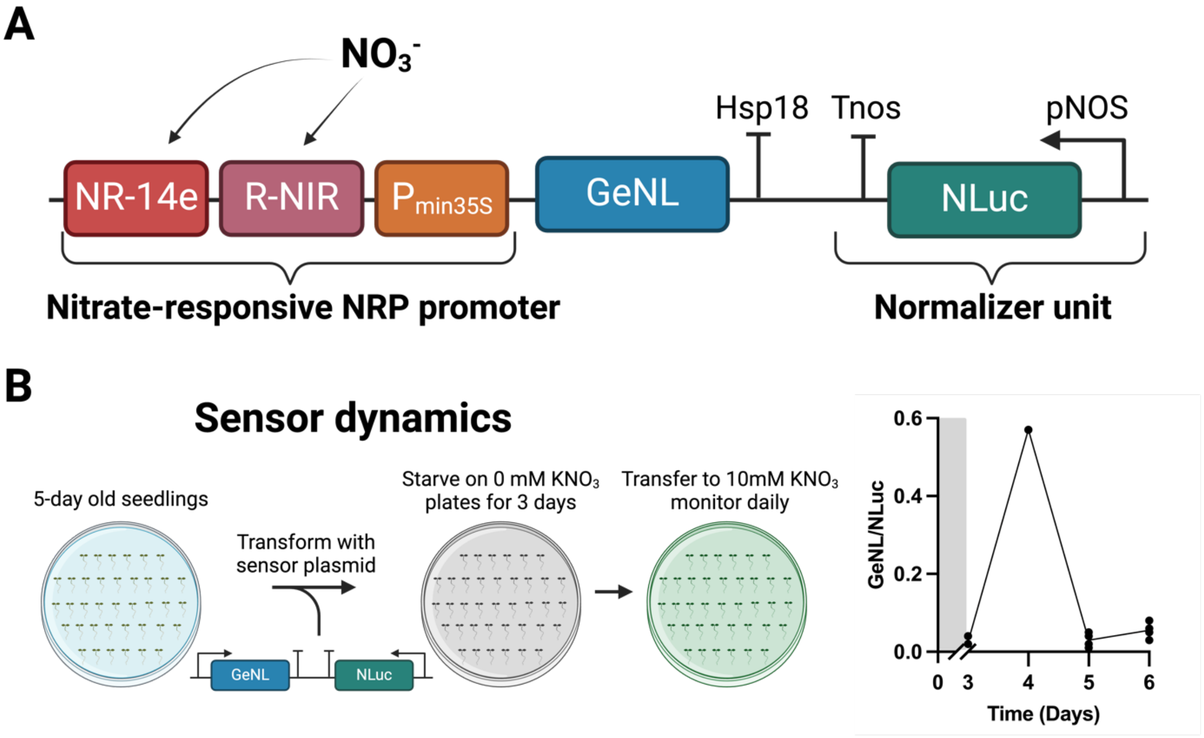


**Supplementary Figure 16. Functional analysis of the Nitrate-Regulated Promoter (NRP) in *S. italica.* (A)** Schematic representation of the construct used. GeNL (Green Enhanced Nano-lantern), NLuc (NanoLuc luciferase), pNOS (nopaline synthase promoter), Tnos (nopaline synthase terminator), and Hsp18 (*Arabidopsis* heat shock protein 18.2 terminator). **(B)** Workflow schematic for testing sensor dynamics and daily luminescence activity of NRP. Each dot represents a biological replicate (4 biological replicates with 5 seedlings per replicate), and the line represents the mean of the replicates. The gray-shaded area indicates the nitrogen starvation period on nitrogen-free plates, used to bring NRP activity to baseline.
